# Population genomics of the inquiline social parasite *Acromyrmex insinuator* and its leaf-cutting ant hosts *A. echinatior* and *A. octospinosus* reveals cryptic differentiation and reduced efficiency of selection in the parasite

**DOI:** 10.64898/2026.08.14.744094

**Authors:** Lukas Schrader, Morten Schiøtt, Rasmus S. Larsen, Mohammed Errbii, Hailin Pan, Qiye Li, Guojie Zhang, Jacobus J. Boomsma

**Affiliations:** Department of Biology, Centre for Social Evolution & Section for Ecology and Evolution, University of Copenhagen, Denmark; Institute for Evolution and Biodiversity, University Münster, Germany; Section for Protein and Enzyme Technology, Department of Biotechnology and Biomedicine, Technical University of Denmark, 2800 Kgs. Lyngby, Denmark; State Key Laboratory of Genome and Multi-omics Technologies, & Shenzhen Key Laboratory of Forensics, BGI Research, Shenzhen 518083, China; College of Life Sciences, University of Chinese Academy of Sciences, Beijing, China; Villum Center for Biodiversity Genomics, Department of Biology, University of Copenhagen, Copenhagen 2100, Denmark; Centre for Evolutionary and Organismal Biology, Women’s Hospital, & Liangzhu Laboratory, School of Medicine, Zhejiang University, Hangzhou 310058, China

**Keywords:** sympatric speciation, population differentiation, gene flow, relaxed selection, genome erosion

## Abstract

Inquiline social parasites usurp colonies of closely related host ants to exploit their social resources. They are almost invariably rare, patchily distributed and difficult to study. Building on almost 25 years of fieldwork in Panama on the social parasite *Acromyrmex insinuator* and its hosts *A. echinatior* and *A. octospinosus*, we use population genomic analyses to test three hypotheses about the evolution of inquiline social parasites: (1) that parasites have extremely reduced effective population sizes, (2) that their genetic variation at coding and non-coding sites shows signatures of eroded adaptive potential, and (3) that they are fully reproductively isolated from their sympatric hosts. We show that the two host species are completely distinct, whereas the parasite has dramatically reduced genetic diversity and effective population size, despite minor ongoing gene flow with its primary host *A. echinatior*. Non-synonymous sites evolve at rates nearly indistinguishable from synonymous sites, indicating a marked reduction in the efficiency of natural selection, consistent with the short evolutionary lifespan of inquiline social parasite lineages. Finally, we uncover clear sub-structure in the *A. insinuator* population, with two genetically distinct lineages co-occurring in the Panama Canal Zone that significantly differ in their likelihood of exploiting the secondary host *A. octospinosus*.

## Introduction

Social parasitism in ants, resulting in mixed colonies of two species, has been known since the 19^th^ century. The phenomenon was extensively covered in *The Origin of Species* (Darwin 1859) and by August Weismann (1893). However, it was Wheeler (1910; 1923; 1928) who for the first time systematically reviewed the global biogeography and natural history of the many convergently evolved lineages of temporary and permanent social parasites, establishing a general understanding that continued in later synthetic reviews (Huxley 1930; Wilson 1971; Hölldobler and Wilson 1990). From Darwin’s days onwards, authors attempted an inclusive understanding that also encompassed various forms of dulosis, the raiding of allospecific colonies to abduct pupae so they can hatch and work for the aggressor’s colony (D’Ettorre and Heinze 2001).

Our current understanding of ant social parasitism, particularly of the non-dulotic inquiline ants, is based on Buschinger’s (1986; 2009) re-interpretation of Emery’s rule (that inquiline social parasites are often sister lineages of their hosts) by conjecturing that many of them arose by sympatric speciation. He also hypothesized that polygynous (multi-queen) host colonies were the ancestral state so that intraspecific queen parasitism could be conceptualized as the first step in a process ultimately leading to reproductive isolation in sympatry. This putative scenario explains why permanent socially parasitic inquilines are restricted to ants, the only lineage of advanced social insects where secondary polygyny is common, resulting in colonies acquiring multiple and only partially related germ-line-like queens that might easily become selected to evolve exploitative traits (Boomsma and Nash 2014). Recent work has corroborated this view, both regarding ancestral polygyny (Boomsma et al. 2014; Dahan and Rabeling 2022) and queen dimorphism and reproductive isolation in sympatry (Leppänen et al. 2016; Wolf and Seppä 2016). New cases of likely sympatric speciation were also identified (Rabeling et al. 2014; Degueldre et al. 2021) while ant lineages with less obligate forms of social parasitism were shown to have evolved allopatrically (Borowiec et al. 2021). Also the remarkable inverse latitudinal diversity gradient of inquiline social parasites (lower species diversity at lower rather than higher latitudes), first suggested by Kutter (1968) and Buschinger (1990), was confirmed to be a real phenomenon, albeit only in the Northern hemisphere (Gray and Rabeling 2023).

Since the 1990s the *Acromyrmex* leaf-cutting ants have become a key genus for studying obligate inquiline parasitism. A highly derived inquiline parasite of South American *Acromyrmex*, so deviant that it was initially described as a separate genus *Pseudoatta*, has been known for more than a century (Gallardo 1916) but discoveries accelerated in recent decades with the identification of *A. insinuator* in Panama (Schultz et al. 1998), *A. ameliae* in Brazil (De Souza et al. 2007), *A. charruanus* in Uruguay (Rabeling et al. 2015), and *A. fowleri* in Brazil (Rabeling et al. 2019). The genus *Acromyrmex* also stands out within the entire clade of the attine fungus-growing ants: inquiline species are unknown from 14 of the 16 genera and only one (sympatrically evolved) case has been documented outside *Acromyrmex* in the genus *Mycocepurus* (Rabeling et al. 2014).

The first comparative genomics study focused on three of the *Acromyrmex* inquilines and their closely related hosts (Schrader et al. 2021). This research identified convergent patterns of general genome erosion due to relaxed selection for ancestral social traits and revealed consistent loss of olfactory receptor genes, a result recently reinforced by a larger comparative analysis across twelve ant subfamilies that also included more attine ant species (Vizueta et al. 2025). These genomic studies indicated that the typically very small and fragmented populations of inquiline social parasites were consistent with observed patterns of gene loss, suggesting that genetic drift could apparently remove or rearrange many ancestral genes without compromising the potential for some allopatric adaptive radiation. The *Acromyrmex* inquilines studied so far invariably have very short coalescence times (range 2.5 - 1 MY) to their free-living host lineages (Schrader et al. 2021), but their genomic erosion does not preclude some potential for adaptive radiation as the highly derived *Pseudoatta* inquiline shares a recent common ancestor with another, less modified inquiline *A. charruanus* (Schrader et al. 2021).

While the last few decades have thus seen major advances in our understanding of ant inquiline biology – taxonomically, phylogenetically and genomically – their consistent rarity continues to largely preclude in-depth studies of the genetic population structure, the ecological habitat characteristics, and the behavioral adaptations that allow inquilines to be efficient exploiters of their host colonies. The only partial exception to this rule is the population of *A. insinuator* in Gamboa and the southeastern Canal Zone in Panama. First collected in Gamboa as a new species in 1993 (Schultz et al. 1998), this parasite turned out not to be quite as rare as usual. It was also amenable to experimental study because its long-lived host colonies could be maintained under climate-controlled laboratory conditions for several years, enabled by the fact that *A. insinuator* queens do not significantly affect the health of their host colony until they take over reproduction. This enabled a series of studies elucidating aspects of life-history and colony founding (Bekkevold and Boomsma 2000; Howe et al. 2021), evolutionarily derived mating system evolution (Baer 2004; Sumner et al. 2004), cuticular chemistry camouflage of parasite workers and queens (Lambardi et al. 2007; Nehring et al. 2015), and the adaptive nursing functions of parasite workers and their reduced disease resistance (Sumner, Hughes, et al. 2003; Sumner, Nash, et al. 2003). Another facilitating factor was that the Copenhagen team maintained its study program of *Acromyrmex* and other Panamanian attine ants for ca. 25 years so that almost yearly collection efforts (usually in in April-May) could accumulate a considerable number of inquiline colonies. These collections allowed us to embark on the present study which is, to our knowledge, the first genome-resequencing study in any inquiline social parasite species, allowing a detailed analysis of the putative causes and implications of genetic differentiation both for the inquiline social parasite and its sympatric free-living host species.

The objectives of the present study were to: (1) Provide conclusive empirical evidence for the expected depletion of genetic diversity in an inquiline ant social parasite; (2) Reconstruct the speciation history of *A. insinuator* and determine to what extent gene flow with the sympatric sister species persists; and (3) Determine how the peculiar demography and life history of the social parasite affects the capacity for adaptive evolution in protein-coding genes. By combining population genomic analyses with demographic modelling, we demonstrate extreme reductions in genetic diversity and concomitant signatures of genome-wide relaxed selection in *A. insinuator*, as well as limited bidirectional gene flow between the sympatric host and parasite populations in and around Gamboa, but not with the secondary host species that was allopatrically colonized.

## Methods

### Sampling, genome re-sequencing, read-mapping and variant calling

Systematic collection of *Acromyrmex* species in and around Gamboa, Panama started in 1993 and continued until 2018. Across 23 annual field campaigns, more than 500 *Acromyrmex* colonies were excavated and, to the extent possible, preserved in freezers in Copenhagen. Many of these colonies were collected with a reproductive queen (sometimes with 2-3 queens; (Nehring et al. 2018)) and were subsequently kept in live culture until they died of natural causes or because of experiments. Winged males and gynes (virgin queens) could be collected for a substantial fraction of the field colonies and from colonies maintained in the lab. The freezer collection thus allowed us to obtain a considerable cross-section of samples from Gamboa for genome sequencing, which we supplemented in 2016 by expanding geographic coverage to include two more remote additional sites in western Panama. We thus obtained a total of 167 colony samples of *Acromyrmex* gynes and queens from three different species and up to three different sites. For *A. echinatior*, the primary host of *A. insinuator* (Schultz et al. 1998), we had 95 samples spanning the sites Cerro Zuela (CAe; n=4), Boca Brava (BAe; n=50), and Gamboa (GAe; n=41). For *A. insinuator*, we had 69 samples, only from Gamboa and the southeastern Panama Canal zone (GAi) because the parasite was not found at the other two locations, and for the outgroup, the secondary host *A. octospinosus*, we included three samples from the Gamboa area (GAo). Detailed information for all samples, collected between 1993 and 2016 and stored in 100% ethanol at -20°C until DNA extraction, is provided in Supplementary Table S1.

Genomic DNA was extracted from individual samples via a conventional salting-out protocol, followed by paired-end library preparation using the Illumina DNA Prep kit and sequencing on an Illumina HiSeq X Ten platform. Extractions had an average yield of 1.47 µg and sequencing generated between 3.3 to 12.9 Gb of data per individual. After adapter trimming with trimmomatic (v0.39), FASTQ files were first converted to unmapped BAM (uBAM) format using Picard (v2.25.2) FastqToSam, after which adapter sequences were marked using Picard MarkIlluminaAdapters, following GATK best-practice recommendations. Mapping was then performed via a three-step pipeline following GATK guidelines for germline short-variant discovery: (1) uBAM files were converted back to FASTQ after which adapter sequences were discarded using Picard SamToFastq; (2) reads were aligned to the reference genome (Vizueta et al. 2025) using BWA-MEM2 (v2.2.1, bwa mem -M) (Li and Durbin 2009); and (3) alignment metadata were restored and adjusted using Picard MergeBamAlignment. The median insert size across samples was ∼231 bp, as assessed with bamtools (Barnett et al. 2011). PCR duplicates were marked and removed from the mapped BAM files using GATK MarkDuplicatesSpark.

Overall mapping statistics (total reads, mapping rate, proper-pair rate, and insert size) were assessed for each sample using bamtools. Per-sample genome-wide coverage was computed using bedtools genomecov (Quinlan and Hall 2010) and further evaluated with Qualimap (Okonechnikov et al. 2016). MultiQC v1.10.1 was used to aggregate and visualise QC metrics across all samples (Ewels et al. 2016). Mapping rates were above 80 % for most samples, with just 12 of the 167 samples having mapping rates of <50 % (Supplementary Figure S1). Fourteen samples (GAi-010, GAi-064, GAi-066, GAi-071, BAe-001, BAe-002, BAe-006, BAe- 007, BAe-009, BAe-010, BAe-011, BAe-012, BAe-025, and CAe-102) where more than 50 % of the genome was covered at less than 9X or where the overall median coverage was below 9x were flagged as poor quality samples and eventually removed (Supplementary Figure S1).

Per-sample coverage bedgraph files were computed and regions exceeding a depth of 80x were extracted as BED intervals for each sample. These per-sample high-coverage intervals were then merged across all samples using bedtools merge. Regions with excess coverage (>80×) in more than 10 individuals (spanning >10 Mb) were compiled into a blacklist BED file for subsequent exclusion during variant filtering, to exclude genomic regions where the assembly might have contained collapsed repeats.

SNP calling followed the GATK v4.2.0.0 best-practice pipeline for germline short-variant discovery (Poplin et al. 2018). Per-sample variant calling was performed using GATK HaplotypeCaller in GVCF mode, after which joint genotyping was carried out with GATK GenotypeGVCFs, producing a single multi-sample Variant Call File (VCF) per scaffold. Per-scaffold VCFs were concatenated into a genome-wide VCF using Picard MergeVcfs and subsequently split into separate SNP-only and indel-only VCFs with GATK SelectVariants.

### Variant filtering and haplotype phasing

Because Variant Quality Score Recalibration (VQSR) requires a large, well-curated set of known variants not available for *A. echinatior*, hard filtering was applied to the SNP call set using GATK VariantFiltration. Variants failing any of the following thresholds were flagged: QD < 2.0, QUAL < 30.0, SOR > 3.0, FS > 60.0, MQ < 40.0, MQRankSum < -5.0, or ReadPosRankSum < -5.0. Only biallelic SNPs passing all filters were retained using GATK SelectVariants.

Missingness-based filtering was performed with VCFtools v0.1.16 (Danecek et al. 2011). In an initial coarse pass, only variants called in at least 50% of individuals and with a minor allele count (MAC) ≥ 3 were retained (--max-missing 0.5 --mac 3). Population-level missingness was then assessed separately for each population (GAe, BAe, CAe, GAi, GAo) using VCFtools -- missing-site. Sites with >10% missing data in any of the three large populations (GAe, BAe and GAi) were excluded. A final, stricter global missingness filter (--max-missing 0.95) was applied subsequently, retaining only variants genotyped in ≥95% of all individuals.

Genomic regions identified as having excess coverage in more than 10 individuals (>80x; see Mapping Quality Control) were removed from the filtered VCF using VCFtools --exclude-bed, masking >10 Mb of potentially misassembled sequence. The final filtered dataset comprised 153 individuals (41 BAe, 3 CAe, 41 GAe, 65 GAi, and 3 GAo) and 1,779,961 biallelic SNPs, after which statistical phasing was performed using SHAPEIT4 v4.1.3 (Delaneau et al. 2012). Genetic map files were constructed per scaffold assuming a uniform recombination rate of 6.2 cM/Mb based on a published linkage map for *A. echinatior* (Sirviö et al. 2006).

### Population structure and cross-species admixture analyses

To visualize genome-wide genetic relationships among all individuals, an identity-by-state (IBS) distance matrix and hierarchical clustering dendrogram were computed using the SNPRelate package in R (Zheng et al. 2012). Prior to distance calculation, SNPs were LD-pruned using *snpgdsLDpruning()* with an r^2^ threshold of 0.9. Hierarchical clustering of the IBS matrix was then performed with *snpgdsHCluster()*, and the resulting tree was cut into groups using *snpgdsCutTree()* with a Z-score threshold of 15 and a permutation test of 5,000 replicates to identify outlier individuals.

Principal component analysis (PCA) with PLINK v1.90 was performed using LD-pruned SNPs (done with PLINK2, --indep-pairwise 50 10 0.1) across all 153 samples (Purcell et al. 2007; Chang et al. 2015). The resulting eigenvectors and eigenvalues were visualized with custom R scripts after which separate PCAs were carried out for the three populations with large sample sizes: Gamboa *A. echinatior* (GAe), Boca Brava *A. echinatior* (BAe) and Gamboa *A. insinuator* (GAi).

Ancestry proportions were estimated using ADMIXTURE v1.3.0 with cross-validation (--cv flag) across a range of K values (Alexander et al. 2009). The LD-pruned SNP dataset for the 153 samples was converted to PLINK BED format using PLINK v1.90 (--make-bed --allow-extra-chr). Models were run for K = 1 to 5, and the optimal K was selected based on the lowest cross-validation error.

Fine-scale population structure and haplotype sharing were investigated using fineSTRUCTURE v4.1.1 (Lawson et al. 2012), which models chromosome painting by ChromoPainter to identify shared haplotype chunks among individuals and cluster them into genetically homogeneous groups. The analysis was run on the final 153-sample SNP dataset. Per-scaffold VCF files were extracted for the 19 chromosomal scaffolds using bcftools view and converted to ChromoPainter phase format using the vcf2cp.pl conversion script. Results were visualised using the accompanying plot.finestructure.R script.

Pairwise genetic differentiation (relative F_ST_ and absolute d_xy_) and nucleotide diversity (π) were calculated for non-overlapping sliding 100 kb windows across the genomes using pixy (Korunes and Samuk 2021). Estimates of d_xy_ and π were corrected for the total number of callable sites per window.

### Demographic modeling with fastsimcoal2

For demographic modeling, we computed folded site-frequency spectra (SFS) for the GAe and GAi samples using easySFS v0.01 (https://github.com/isaacovercast/easySFS). The SFSs were then downprojected, maximising the number of segregating sites retained. To infer demographic history and test for gene flow between the Gamboa host ant and social parasite populations (of which the latter turned out to have two genetically distinct morphs, which we named Ai-left and Ai-right), we fitted five demographic models of increasing complexity to a 2D folded joint SFS (GAe and GAi) and four models to a 3D folded joint SFS (GAe, GAi-left and GAi-right) using fastsimcoal2 v2.8 (Excoffier et al. 2013), assuming a neutral mutation rate of 3.6e-9 (an estimate for bumble bees from Liu et al. (2017) as direct estimates for ants are lacking). The five two-population models tested were: (1) Strict isolation (SI): no gene flow after the two species split up, modelled with 4 parameters; (2) Isolation with symmetric migration throughout (IM), 6 parameters; (3) Isolation with asymmetric migration throughout (IM_asym), 6 parameters; (4) Ancient migration only, i.e., gene flow ceased well before present (AM), 7 parameters; and (5) Secondary contact, i.e., isolation followed by recent gene flow (SC), 7 parameters. The four three population models tested were: (1) Strict isolation (SI): no gene flow between any of the populations, 7 parameters; (2) Isolation with ancestral migration between GAe and the GAi ancestral population (IM3_ANC), 8 parameters; (3) Isolation with migration between GAe and the contemporary GAi populations GAi-left and GAi-right (IM3_REC), 9 parameters; and (4) Isolation with continuous migration between GAe and the GAi ancestral population and the contemporary GAi populations GAi-left and GAi-right (IM3_CONT), 10 parameters.

All five two population models shared a common population tree in which GAe (pop 0) and GAi (pop 1) diverged from an ancestral population. In the three population models, GAe and GAi diverged jointly from an ancestral population and GAi secondarily split into GAi-left and GAi-right. Effective population sizes of GAe and GAi were parameterized in all models while using a dynamic parameter bound to incorporate the constraint that N_POP1 > N_POP2, based on both the present genetic diversity analyses and those by Schrader et al. (2021).

Each model was run for 100 independent optimization replicates, each using 1,000,000 coalescent simulations per parameter proposal and 50 ECM optimization cycles. The replicate with the highest composite log-likelihood (MaxEstLhood) was retained as the best run for each model. Model selection was performed using the Akaike Information Criterion. To obtain confidence intervals for parameter estimates produced by the best-fitting model, we performed non-parametric block bootstrapping. To this end, the VCF was split into 100 equal-sized genomic blocks so 100 bootstrap replicates could be generated by resampling. A new folded SFS was then computed for each bootstrap replicate again using easySFS with the same projection sizes, after which the best-fitting model was re-fitted to each replicate using 20 independent optimization runs per replicate and 50 ECM cycles. The 2.5th and 97.5th percentiles of the resulting parameter distributions were then used as 95% confidence intervals.

### Demographic modeling with smc++

Effective population size through time was inferred separately for the six populations (BAe, CAe, GAe, GAi-left, GAi-right, GAo) with SMC++ v1.15.4. Regions not callable in ≥95% of the 153 samples were masked, and smc++ inputs were generated for the 19 largest scaffolds of the assembly, using three randomly selected distinguished individuals per population. Histories were estimated under a cubic spline assuming µ = 3.6e-9 (see above). Uncertainty was assessed by block bootstrapping: 1.5 Mb chunks were resampled with replacement into 20 pseudo-chromosomes of 10 chunks, written as separate files, and 100 replicates per population were refitted.

### Demographic modeling with MSMC2

Effective population sizes and relative cross-coalescence over time were reconstructed with MSMC2 using phased SNP data from four diploid individuals, i.e., eight phased haplotypes. Chromosome-specific MSMC2 multihetsep files were generated using a callable-genome mask derived from the complement of regions classified as uncallable in at least 5% of samples. Genomic uncertainty was assessed using 134 bootstrap replicates generated by resampling 5-Mb genomic blocks into 19 pseudochromosomes containing three blocks each, corresponding to approximately 285 Mb per bootstrap genome. For each replicate, MSMC2 was run separately for each population, and 64 times for each between-population haplotype comparison. Within- and between-population results were subsequently combined using combineCrossCoal.py. Effective population size was retrieved from the MSMC2 coalescence parameter as N_e_ = 1/(2μλ), and relative cross-coalescence was calculated as 2 x λ_01_/(λ_00_ + λ_11_). Final demographic trajectories were summarized across the 134 genomic bootstrap replicates using the median and empirical 2.5th and 97.5th percentiles.

### McDonald-Kreitman analysis of protein-coding evolution

To test for differences in the mode and efficacy of selection on protein-coding sequences among the three focal populations (GAi, GAe, BAe), we performed McDonald-Kreitman (MK) tests using the *A. octospinosus* population (GAo) as outgroup. Sites were classified as 0-fold degenerate (every change nonsynonymous) or 4-fold degenerate (every change synonymous) using degenotate (https://github.com/harvardinformatics/degenotate). Variants from the joint VCF were partitioned by these degeneracy classes, so that counts of polymorphic (pN, pS) and fixed (dN, dS) sites were tabulated per gene for each focal population separately, treating GAo as the outgroup for divergence calling. MK summary statistics were computed for each population by pooling counts across all genes with ≥1 polymorphic sites and ≥1 fixed differences: the polymorphism ratio (pN/pS), the divergence ratio (dN/dS), the neutrality index (NI = (pN/pS)/(dN/dS)), and the fraction of substitutions driven by positive selection (α = 1 − NI (Smith and Eyre-Walker 2002)). Uncertainty was quantified by non-parametric bootstrapping (2,000 replicates), resampling genes with replacement, with 95 % confidence intervals derived from the 2.5th and 97.5th percentiles of the bootstrap distribution. Pairwise differences between populations were then tested by computing the bootstrap distribution of paired statistical differences and reported as two-sided percentile p-values, with Benjamini-Hochberg correction across the three population pairs. We retained only genes for which all four counts (pN, pS, dN, dS) were reported by degenotate, and for which at least one polymorphism and one substitution was recorded. CAe was excluded from the MK analysis because of the small sample size (n = 3).

### Site Frequency Spectrum Construction

For DFE-inference with polyDFE, nonsynonymous and synonymous variants were annotated and extracted with SnpEff (v5.1) with a custom gene model database derived from the most recent *A. echinatior* genome annotation (Vizueta et al. 2025). Ancestral allelic states were inferred using est-sfs v2.04 (Keightley and Jackson 2018) with *A. octospinosus* as outgroup; however, due to considerable polarization error causing an excess of high-frequency derived alleles (affecting >20% of sites based on diagnostic checks), we opted for using the folded site-frequency spectrum for subsequent analyses which is independent of the ancestral allele states (see below).

We generated folded site-frequency spectra (SFS) for both synonymous and non-synonymous codon sites using easySFS with projection to 56 haplotypes per population (80% of GAi samples, 67% of GAe samples). The final dataset comprised 1,530 segregating synonymous sites and 989 segregating missense sites in GAi (projected to 56 haplotypes), and 2,696 segregating synonymous sites and 1,548 segregating missense sites in GAe. To quantify divergence, we then computed the number of fixed differences (sites fixed for alternative alleles between lineages) for missense (*D*_n_) and synonymous (*D*_s_) sites. Allele frequencies were calculated separately for GAi, GAe, and the *A. octospinosus* outgroup (GAo; *n* = 3 diploid samples) using BCFtools +fill-tags. For outgroup-rooted divergence, we identified sites where GAi (or GAe) was fixed for one allele and GAo was fixed for the other allele. We then quantified GAi vs. GAo divergence (*D*n = 18,170; *D*s = 33,127) and GAe vs. GAo divergence (*D*n = 15,356; *D*s = 28,366), yielding *D*n/*D*s ratios of 0.548 and 0.541, respectively.

### Distribution of Fitness Effects (DFE)

We inferred the distribution of fitness effects (DFE) for new nonsynonymous mutations using polyDFE v2.0 (Tataru et al. 2017; Tataru and Bataillon 2019). PolyDFE jointly models the folded SFS and divergence data to estimate the shape of the DFE assuming a gamma distribution with parameters *S*_d_ (mean deleterious selection coefficient, scaled by 2*N*e) and a shape parameter *b*. We compared two demographic-DFE models: Model B, which assumes all nonsynonymous mutations are deleterious or neutral (*p_b_* = 0, where *p*_b_ is the proportion of beneficial mutations), and Model C, which allows for a class of beneficial mutations with mean selection coefficient *S*_b_. For each population and model, we performed 20 independent optimization runs with random starting parameters (S_d_ ∼ log-uniform[−1, −1000], *b* ∼ uniform[0.1, 1.0], *p*_b_ ∼ uniform[0.0, 0.5], *S*_b_ ∼ log-uniform[1, 1000]), and selected the run with the highest log-likelihood. Critically, we fixed the ancestral-allele misidentification rate parameter (ε_an_) at 0, as our folded SFS does not use ancestral state information. Demographic nuisance parameters (*r*2…*r*56, representing deviations from the standard neutral coalescent) were initialized from the observed synonymous SFS and estimated during optimization.

The proportion of substitutions driven by positive selection (α_div_) was estimated from the best-fit Model C parameters following Eyre-Walker & Keightley (2009). Briefly, α represents the fraction of nonsynonymous substitutions caused by beneficial mutations fixed due to positive selection rather than genetic drift. We estimated α using the divergence-based method implemented in polyDFE, which compares the observed ratio of nonsynonymous to synonymous divergence (*D*_n_/*D*_s_) with expectations based on neutrality and exclusive purifying selection while accounting for slightly deleterious mutations that fix due to drift. Confidence intervals (95%) for all DFE parameters and α values were obtained through parametric bootstrapping with 200 replicates. For each of these replicates, we resampled the SFS from the multinomial distribution defined by the best-fit Model C parameters and re-estimated all parameters. Model comparison between Model B and Model C was performed using likelihood ratio tests (LRT) with 2 degrees of freedom (for *p*_b_ and *S*_b_).

## Results

### Population genomic sequencing, SNP diversity, and population structure

We sequenced and analysed whole genomes from queens or gynes of the inquiline social parasite *A. insinuator* and of its two Panamanian hosts species *A. echinatior* and *A. octospinosus* using short-read sequencing. After read mapping against the chromosome-level *A. echinatior* reference genome (Vizueta et al. 2025) and stringent variant filtering (see Methods), the final dataset comprised 153 gyne/queen individuals (3 GAo, 41 BAe, 3 CAe, 41 GAe, 65 GAi, Supplementary Figure 1; Supplementary Table 1) and 1,779,961 biallelic SNPs. In spite of only three colonies having been included, *A. octospinosus* showed the highest total SNP count (915,617), implying this species was genetically most distinctively different from the *A. echinatior* reference genome. A total of 255,700 (28.0%) of these SNPs were polymorphic across the *A. octospinosus* individuals. Within the *A. echinatior* gene pool, the Boca Brava population had the highest number of SNPs (n_total_ = 646,294; n_poly_ = 627,544 (97.1%)), followed by Gamboa (n_total_ = 443,718; n_poly_ = 443,681 (99.9%)), whereas the Cerro Zuela population had fewest SNPs (n_total_ = 337,614; n_poly_ = 255,674 (75.7%)), most likely reflecting the small sample size (n = 3 colonies) rather than reduced genetic diversity (see below). The *A. insinuator* population in Gamboa had substantially more SNPs (n_total_ = 560,943) than any of the three *A. echinatior* populations, reflecting approximately 1 MYA of independent evolution since the two species diverged (Schrader et al. 2021). Of these *A. insinuator* SNPs 126,227 (22.5%) were polymorphic. While SNP counts and polymorphisms are affected by sample sizes and population structure, these overall results appear consistent with Panamanian *A. octospinosus* being a continuously distributed rain forest species with relatively higher effective population sizes and with *A. echinatior* being a more patchily distributed species characteristic of open/disturbed habitat. This would then match the earlier result that *A. octospinosus* also has higher heterozygosities at microsatellite marker loci (Ortius-Lechner et al. 2000). In this perspective the SNP count of the social parasite *A. insinuator* is remarkably high.

Hierarchical clustering in the pairwise identity-by-state (IBS) distance matrix resolved three strongly supported groups corresponding hierarchically to species and sampling locality within species (Figure 1A). The deepest split separated the three *A. octospinosus* individuals from all other samples, consistent with their distinct species status (Schultz et al. 1998) and the greater phylogenetic distance of *A. octospinosus* from the other two species (see also Vizueta et al. (2025), which estimated the two free-living Panamanian *Acromyrmex* species to have separated 3.2 MYA). The second major split separated the 65 *A. insinuator* individuals from the three *A. echinatior* populations, confirming strong genetic divergence between the social parasite and its primary host despite their sympatric occurrence in Gamboa and in line with an earlier estimate of their divergence time of ∼1 MYA (Schrader et al. 2021). Within *A. echinatior*, the Gamboa (GAe) and Boca Brava (BAe) populations formed well-differentiated sister clusters but with a much shallower divergence than the higher-level segregation between *A. echinatior* and *A. octospinosus*. The three Cerro Zuela individuals/colonies (CAe) clustered at the base of the BAe clade, suggesting higher overall genomic affinity with Boca Brava than with the geographically closer Gamboa population. However, one Boca Brava individual (BAe-107) fell unambiguously within the GAe clade with IBS distances comparable to within-GAe comparisons, suggesting either a recent dispersal event of a Gamboa-gyne to Boca Brava or an inadvertent sample swap. This individual was re-assigned to the GAe population in all subsequent analyses.

**Figure 1:**
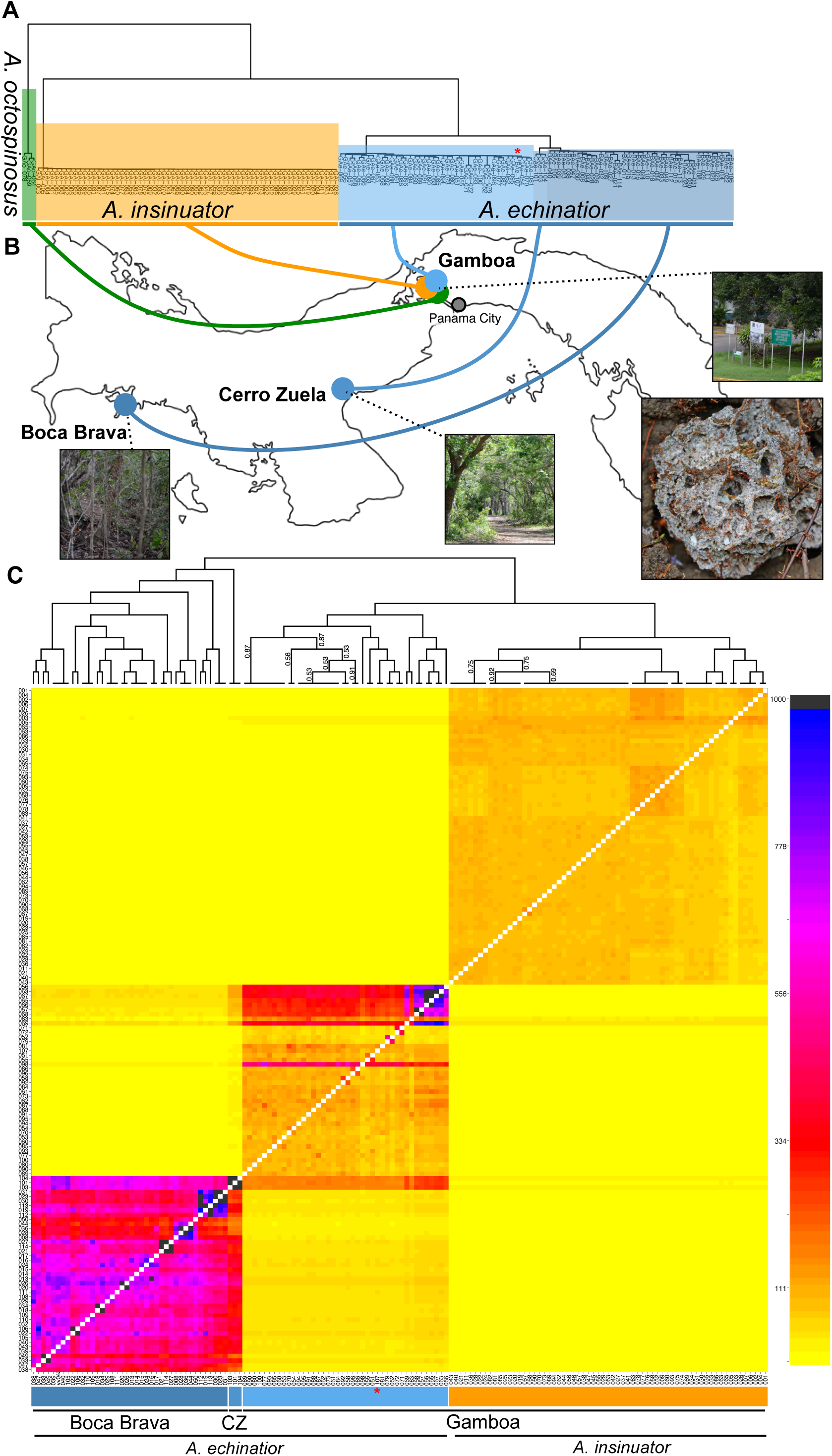
Sampling localities, population differentiation and population structure of *A. echinatior*, *A. octospinosus* and *A. insinuator* populations in Panama. (**A**) IBS-based dendrogram of 153 samples (*A. echinatior*: 41 Boca Brava (BAe; dark blue), 41 Gamboa (GAe; blue), 3 Cerro Zuela (CAe; light blue), 65 *A. insinuator* (GAi; orange), and 3 *A. octospinosus* (GAo; green). Habitat images of sampling localities and a picture of a typical *Acromyrmex* fungus garden are also provided. (**B**) Heatmap of colony-level co-ancestry across the sister species *A. echinatior* and *A. insinuator* generated with FineStructure. The dendrogram represents hierarchical clustering of individuals based on total genome-wide haplotype sharing inferred by ChromoPainter, with branch lengths reflecting the degree of genomic similarity between individuals or groups. One sample of *A. echinatior* from Boca Brava (BAe-107) clustered within the Gamboa population (red asterisk) consistent with a recent dispersal event or sample swap. Numbers in the dendrogram indicate bipartition certainty (proportion of MCMC iterations supporting each split) when certainty values were < 1. Color intensity illustrates the degree of haplotype sharing between putative haplotype donors (columns) and recipients (rows).

PCA on the LD-pruned SNPs across all 153 samples recapitulated the three-species separation (Supplementary Figure 2). The first two PCs jointly explained 78.6% of total variance (PC1 = 43.4%, PC2 = 35.2%), with each species occupying a distinct, well-separated cluster in PC space. ADMIXTURE analysis of the LD-pruned SNPs supported K = 4 ancestral populations as the best-fitting model (CV error = 0.125; Supplementary Figure 3), separating the 153 samples into four groups corresponding to *A. echinatior* from Gamboa, *A. echinatior* from Boca Brava, *A. insinuator* from Gamboa, and *A. octospinosus* from Gamboa. In this analysis, the Cerro Zuela individuals were assigned predominantly to the Boca Brava ancestry component, consistent with their placement in the dendrogram (Figure 1A). ChromoPainter/fineSTRUCTURE analysis of genome-wide haplotype sharing further corroborated the species-level separations and revealed finer details of gene flow at the species boundaries (Figure 1B). The co-ancestry heatmap and hierarchical clustering revealed clear genomic separation between *A. insinuator* and *A. echinatior*, but with evidence of some haplotype sharing between the sympatric Gamboa samples (GAe and GAi) and more subtly between *A. insinuator* and the Cerro Zuela population of *A. echinatior* (CAe). Consistent with localized gene flow in sympatry, *A. insinuator* and *A. echinatior* in Gamboa clustered more closely to one another than did the allopatric Boca Brava and Gamboa populations of *A. echinatior* (BAe and GAe) (see dendrogram in Figure 1B). This suggests that ongoing haplotype exchange between the two species at this site exceeds the cumulative haplotype sharing between the allopatric populations of *A. echinatior* and would be consistent with the considerable population viscosity in the GAe population that we document below. Within *A. echinatior*, the Cerro Zuela samples grouped once more with the Boca Brava samples while showing elevated haplotype sharing with Gamboa relative to the rest of the Boca Brava cluster. Asymmetry in haplotype donor-recipient relationships between *A. echinatior* populations suggests a net excess of gene flow from Boca Brava towards Gamboa rather than in the reverse direction.

### Population sub-structure

We computed population-specific Principal Components to identify signatures of within-population substructure for the three populations with sufficient sample sizes: BAe, GAe, and GAi. The *A. echinatior* populations in Boca Brave (BAe, Figure 2A) and in Gamboa (GAe, Figure 2B) each showed one main cluster containing most individuals. In contrast, a strong signature of population sub-structuring with two well defined clusters separated primarily along PC1 (named GAi-left and GAi-right) occurred in the Gamboa social parasite population (Figure 2C). Analyses of our field notebook collection records revealed that GAi-right individuals were collected predominantly before the year 2000 in the village of Gamboa and at not previously sampled locations towards Panama City after 2000. In contrast, the GAi-left individuals were mostly collected in the village of Gamboa after the turn of the century (Figure 2D, Fisher test, p<0.0001, odds ratio = 32.19). ChromoPainter/fineSTRUCTURE analyses restricted to the GAi samples provided further evidence for population sub-structuring in the inquiline parasite population (Figure 2E). Using ADMIXTURE, three of the four early *A. insinuator* samples with low sequencing quality could also be assigned to GAi-right (GAi-064 from 2009 and GAi-066 from 2010) and GAi-left (GAi-071 from 2010), respectively (Supplementary Figure S4; Supplementary Table 1), which hardly affected the correlation of Figure 2D (p<0.0001, odds ratio = 22.52).

**Figure 2:**
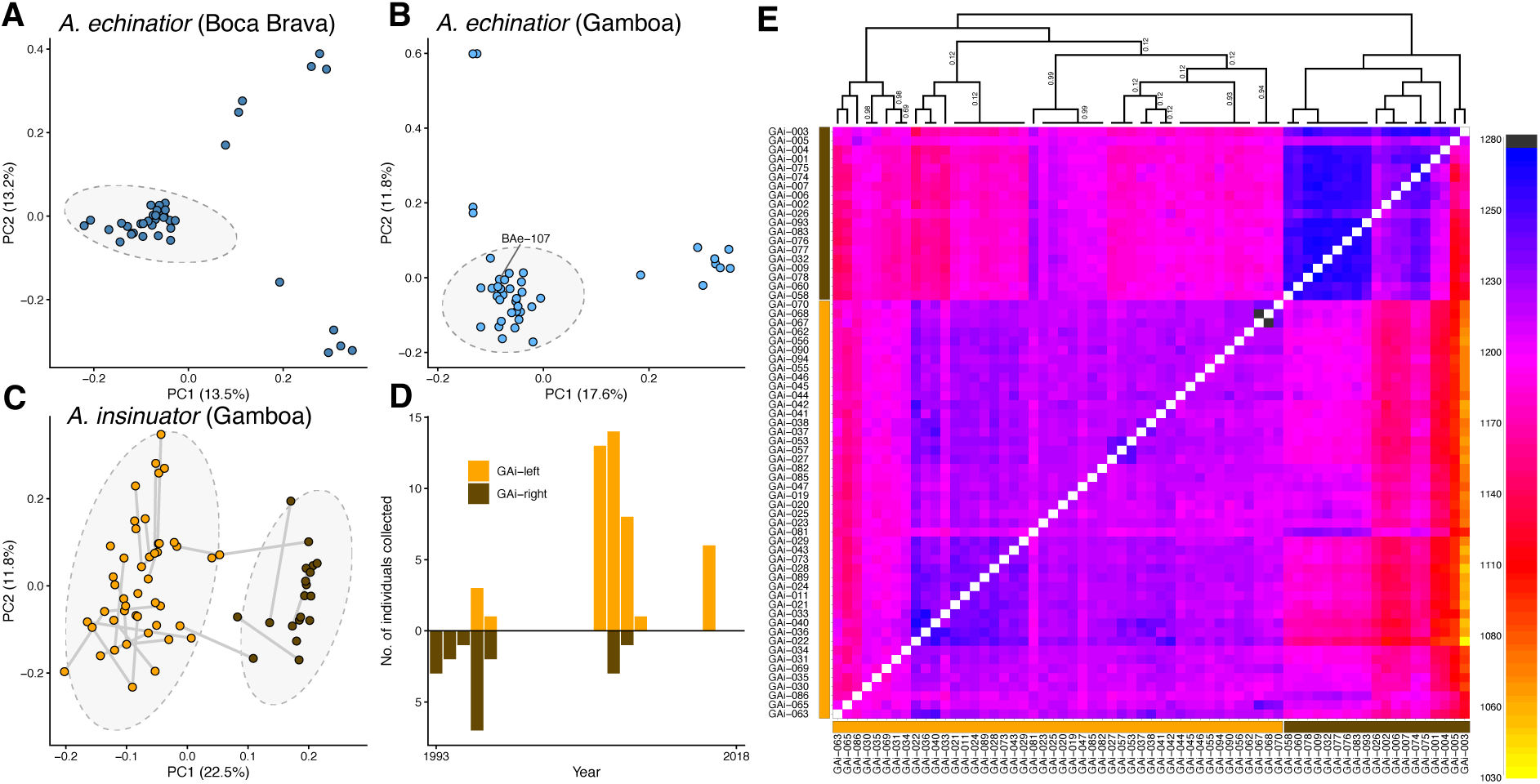
Population substructure and differentiation. (**A-C)** Population-specific PCAs for *A. echinatior* from Boca Brava (**A**, BAe), *A. echinatior* from Gamboa (**B**, GAe), and *A. insinuator* from Gamboa (**C**, GAi). Grey ellipses highlight either the single main cluster in the two *A. echinatior* populations (**A**, **B**) or the two clusters GAi-left and GAi-right sub-structuring the *A. insinuator* population in Gamboa and the southeastern Panama Canal range into two distinct demes (**C**). **(D)** Collection history of individuals from the two *A. insinuator* demes GAi-left (orange) and GAi-right (brown) in Gamboa between 1993 and 2016, showing that GAi-right was predominantly collected before and GAi-left after the year 2000 **(E)** Heatmap of co-ancestry across the 69 *A. insinuator* samples identified with fineSTRUCTURE, with GAi-left (bottom-left) and GAi-right (top-right) appearing as two distinct clusters of intensified haplotype-sharing. The dendrogram represents hierarchical clustering of individuals based on total genome-wide haplotype sharing inferred by ChromoPainter, with branch lengths reflecting the degree of genomic similarity between individuals or groups. Numbers in the dendrogram indicate bipartition certainty (proportion of MCMC iterations supporting each split) when certainty values were < 1.

Mapping of the *A. insinuator* sampling localities in (Figure 3A) and around (Figure 3B) Gamboa revealed no clear geographic signal to explain the two genetic lineages identified by our population structure analyses (Figure 2C). The sampled individuals, collected predominantly from *A. echinatior* host colonies but over time increasingly also from *A. octospinosus* colonies (or sometimes as lone wandering queens), were distributed rather evenly across the total sampling area, consistent with overall sympatry. However, the *A. octospinosus* nests infested with parasite queens were invariably from the forested (darker green) area in Gamboa (or from the edges of it), whereas *A. echinatior* was found outside the forest or along the more sunlit banks of the Gamboa pond (Laguna de Gamboa) inside that forest area (see Figure 3A), which extended further North in the 1990s and 2000s than this more recent map indicates. In two instances, *A. insinuator* individuals from both demes were obtained from the same field-sampled host colony (Ao054, Ae413), indicating that the two inquiline lineages can co-occur within a single host nest, both of the primary host *A. echinatior* and of the secondary host *A. octospinosus* (Figure 3A,B).

**Figure 3:**
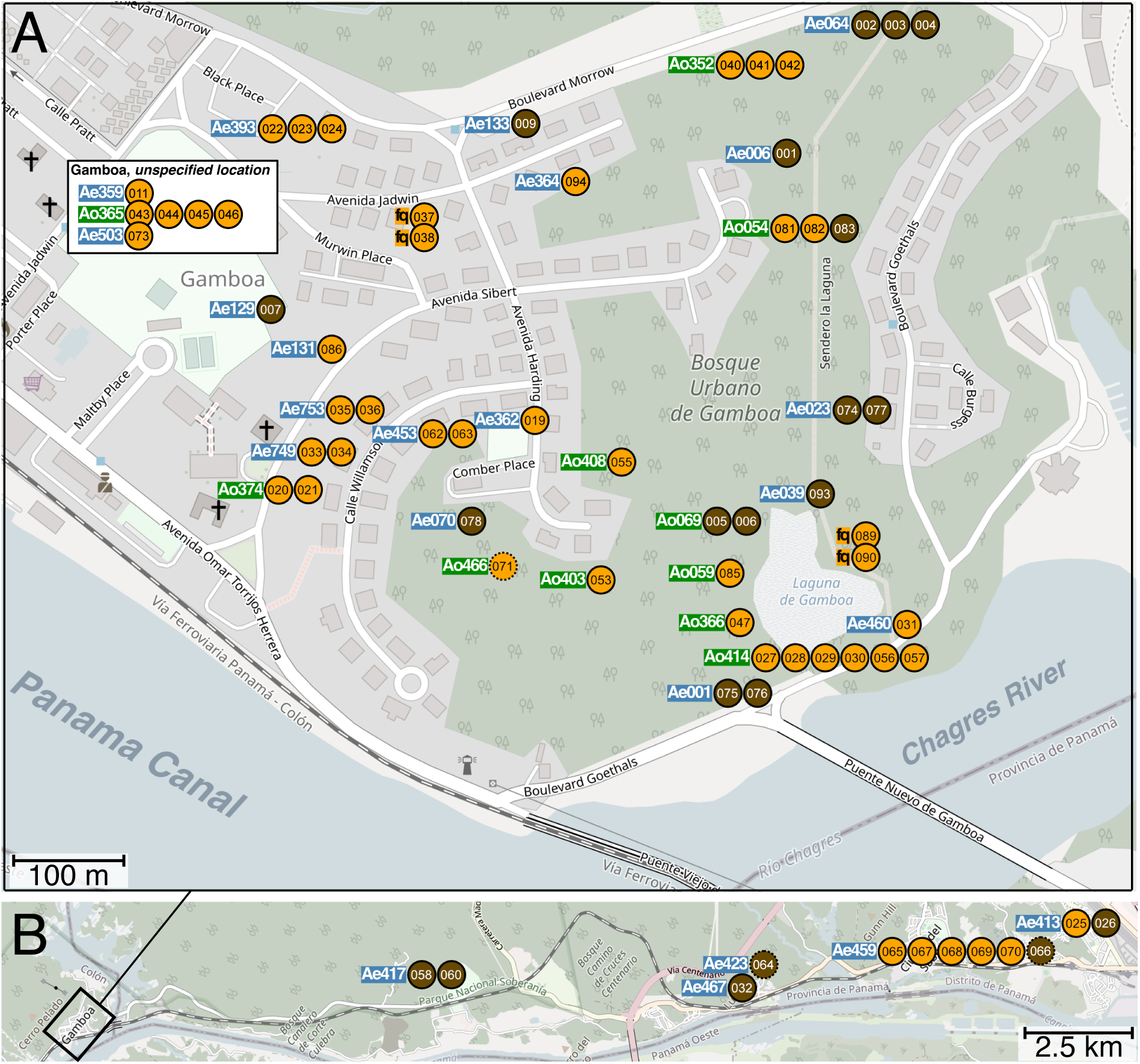
Approximate collection locations of the sequenced *A. insinuator* (GAi) specimens in Gamboa, Panama, and along the Panama Canal. Each circle depicts an individual belonging to the left deme (orange) or the right deme (brown) (Figure 2C) sampled in Gamboa **(A)** or along the Panama Canal towards Panama City, a geographic extension we considered not to be isolated from Gamboa while the samples were collected **(B)**. Rows of numbered circles are *A. insinuator* queens/gynes collected from the same host colony (acronyms in blue for *A. echinatior* and in green for *A. octospinosus* as in Figure 1A). As far as habitat-information was recorded, *A. echinatior* was collected from the grey areas (disturbed and at least half open road verge and garden habitat) while *A. octospinosus* was collected from the wooded (green) areas (smaller wooded patches are not marked in green on the map). Host colonies with IDs up to Ae133 were collected between 1993 and 2000, while those with higher IDs were collected between 2008 and 2016. Lone wandering founding queen IDs (fq) refer to collections in 2009 (sample 89 and 90) and 2016 (sample 37 and 38). Low-quality genome samples (064, 066, 071) are shown with dashed rather than solid circles. See Supplementary Table 1 for further collection details.

Further scrutiny of the field notebooks revealed that for the GAi-right deme (Figure 2C), 16 individuals were collected from *A. echinatior* colonies and only three from *A. octospinosus* colonies, whereas this ratio was even (21/21) for the GAi-left deme, a difference that was significant (p < 0.013; Fisher’s exact test, odds ratio = 0.19). Including the three low-quality samples, these ratios changed to 18/3 for GAi-right and 21/22 for GAi-left (p < 0.006, odds ratio =0.16). Furthermore, after the year 2000, GAi-right individuals were no longer collected in the village of Gamboa but at sites 5-20 km away in southeastern direction towards Panama City (Figure 3B: four parasite queens from *A. echinatior* host colonies in 2009 (Ae413, Ae417) and 2010 (Ae467); six including the low-quality samples). Conversely, the few individuals of the GAi-left deme collected before 2000 were predominantly from *A. octospinosus* host colonies (3 out of 4) and only became common in *A. echinatior* host colonies in the sampling years after 2000 (20 out of 36; 22 out of 38 when including low-quality samples).

When combining the information from these maps (Figure 3A,B) with the GAe genome dispersion patterns obtained by PCA (Figure 2B), it appeared that the eight outliers towards the right along the 1^st^ PC axis of Figure 2B mirror-imaged the increasing distance from Gamboa village across a distance of up to 25 km (Figure 3B), with a correlation between PC1 score and geographical distance from Gamboa village of ρ = 0.91 (p = 0.0005, Spearman rank correlation). This implies a remarkable degree of population viscosity for the GAe population sensu lato across distances an order of magnitude less than the distances from Gamboa to the remote *A. echinatior* populations at Cerra Zuela and Boca Brava (Figure 1A). These eight colonies are also visibly distinct as the blue square in the center of Figure 1B. In contrast, the 4 outliers along the PC2 axis in Figure 2B could not be related to any geographic pattern.

### Nucleotide diversity and differentiation between populations

We next compared genetic diversity and differentiation between different species, populations and demes, calculating nucleotide diversity (π) for each (sub-)population and pairwise relative (F_st_) and absolute divergence (d_xy_) between individual pairs across (sub-)populations in 200 kb windows along the reference genome (Figure 4). Nucleotide diversity (π) differed markedly between socially parasitic *A. insinuator* (both demes combined) and the free-living species (Figure 4A). All three *A. echinatior* populations (BAe, CAe, GAe) as well as *A. octospinosus* showed similar genome-wide diversity (median log10(π) ≈ -4.2 in GAo to -4.1 in BAe). In stark contrast, *A. insinuator* exhibited dramatically reduced genome-wide nucleotide diversity, with a median π approximately 1.5 orders of magnitude lower than in its host species (median log10(π) ≈ -5.6).

**Figure 4:**
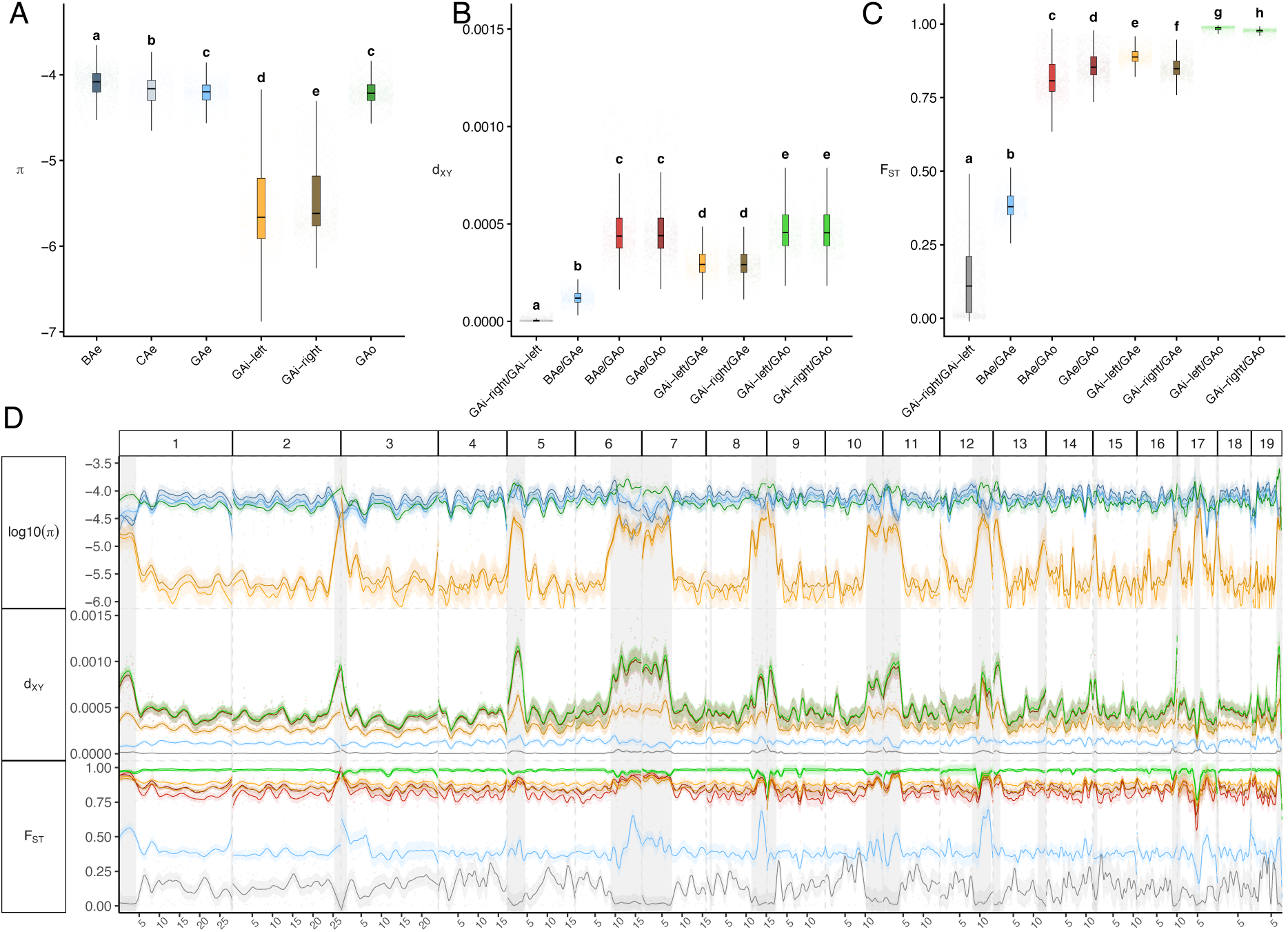
Genome-wide nucleotide diversity and differentiation between Panamanian *Acromyrmex* species and their (sub-)populations against the chromosome-resolved *A. echinatior* reference genome from Gamboa. (**A**) Nucleotide diversity (log10(π)) for *A. echinatior* from Boca Brava (BAe), Cerro Zuela (CAe), and Gamboa (GAe), the two demes of *A. insinuator* from Gamboa (GAi-left and GAi-right) and *A. octospinosus* (GAo) obtained from 200 kb sliding windows along the 19 chromosomes of *A. echinatior*. While the populations of *A. echinatior* and *A. octospinosus* exhibit very little overall difference in nucleotide diversity, the inquiline social parasite *A. insinuator* has dramatically reduced nucleotide diversity (for both clusters plotted in Figure 2C) compared to its primary (*A. echinatior*) and secondary (*A. octospinosus*) host species. **(B)** Genetic differentiation expressed as absolute divergence (d_xy_) obtained from the same sliding windows, showing very low values for the two *A. insinuator* demes compared to the *A. echinatior* populations, intermediate values for the *A. insinuator* demes relative to their primary host *A. echinatior* and the highest values for all comparisons with *A. octospinosus*. **(C)** Genetic differentiation expressed as relative divergence (F_ST_) obtained from the same sliding windows. As expected, the populations of *A. echinatior* from Gamboa and Boca Brava (blue) were only weakly differentiated (median d_xy_ ≈ 1.7e-4, median F_ST_ ≈ 0.4, but the differentiation between *A. echinatior* and A. *octospinosus* (red; GAe/GAo and BAe/GAo) and between *A. echinatior* and *A. insinuator* (orange; GA-left cluster/GAe and GAi-right cluster/GAe) was much stronger (median d_xy_ > 6e-4, median F_st_ > 0.8). The strongest differentiation occurred between *A. insinuator* and *A. octospinosus* (median d_xy_ ∼6.5e-4, median F_ST_ ∼1). Different letters indicate statistically significant differences between groups (p<0.05, pairwise Wilcoxon rank-sum tests, Bonferroni-corrected). (**D**) Overall, genome scans revealed that large-scale deviations in log10(π), d_xy_ and F_ST_ (color coding as in the A-C panels) coincided with regions enriched for transposable elements in the genome of *A. echinatior* (grey background highlights) but without clear signatures of selective sweeps or introgressions between species and/or populations.

Pairwise d_xy_ and F_ST_ estimates confirmed the species-level boundaries inferred from the population structure analyses (Figure 4B,C). Differentiation between Gamboa *A. echinatior* and each of the two *A. insinuator* demes was high (avg. d_xy_ ≈ 2.9e-4, avg. F_ST_ ≈ 0.85) and there was no obvious asymmetry in differentiation of the two GAi demes from *A. echinatior* (GAe), so differential gene flow from the host is unlikely to explain GAi population structure. Between *A. echinatior* and *A. octospinosus,* d_xy_ was even higher (avg. d_xy_ ≈ 4.4e-4, avg. F_ST_ ≈ 0.85) consistent with a considerably deeper phylogenetic split between these two reproductively isolated species (Schultz et al. 1998; Vizueta et al. 2025). We found the strongest differentiation between sympatric *A. insinuator* (GAi) and *A. octospinosus* (GAo) individuals (avg. d_xy_ ≈ 4.6e-4, avg. F_ST_ ≈ 0.98), consistent with no gene flow between the parasite and its secondary host which it must have colonized later than the primary host from which it evolved. In contrast and as expected for conspecific populations, d_xy_ and F_ST_ between the two *A. echinatior* populations (GAe and BAe, mean d_xy_ ≈ 1.2e-4, mean F_ST_ ≈ 0.38) and particularly between the two sympatric *A. insinuator* demes were substantially lower (mean d_xy_ ≈ 2.9e-6, median F_ST_ ≈ 0.11). However, the relatively high F_ST_ value of 0.38 obtained when comparing the GAe and BAe populations seems reasonable when extrapolating the population viscosity estimates over distances up to ca 25 km within the GAe population (Figure 2B; Figure 3B).

Chromosome-level variation in nucleotide diversity (π) and differentiation (d_xy_ and Fst) at the beginning and the end of chromosomes often coincided with putatively pericentromeric accumulations of transposable element identified in the *A. echinatior* reference genome (grey background highlights in Figure 4D) (Schrader et al. 2026). Similar effects on differentiation and genetic diversity were previously described in population genomic analyses of the ant *Cardiocondyla obscurior* (Errbii et al. 2021; Errbii et al. 2024).

Demographic modelling with FastSimCoal2, based on a down-projected folded 2D site-frequency spectrum of the GAe and GAi samples, showed the highest support for a secondary-contact (SC) demographic model (Figure 5A, Supplementary Figure 5). Median estimates derived from parametric block-bootstrapping placed the divergence of the two species ca. 600,000 generations ago (i.e. 1.2 MYA, assuming a generation time of two years), broadly consistent with a previously inferred estimate of ∼0.9 MY divergence (Schrader et al., 2021). This best-fitting, secondary-contact model inferred complete reproductive isolation following speciation, with limited gene flow resuming during the last 700-800 generations and proceeding 2.5 times as efficiently from host to parasite than in the reverse direction (Figure 5A). The contemporary N_e_ of the parasite *A. insinuator* (GAi) was estimated to be 26 k, an order of magnitude below that of the *A. echinatior* host population in Gamboa (N_e_ GAe 271 k) (Figure 5A).

**Figure 5:**
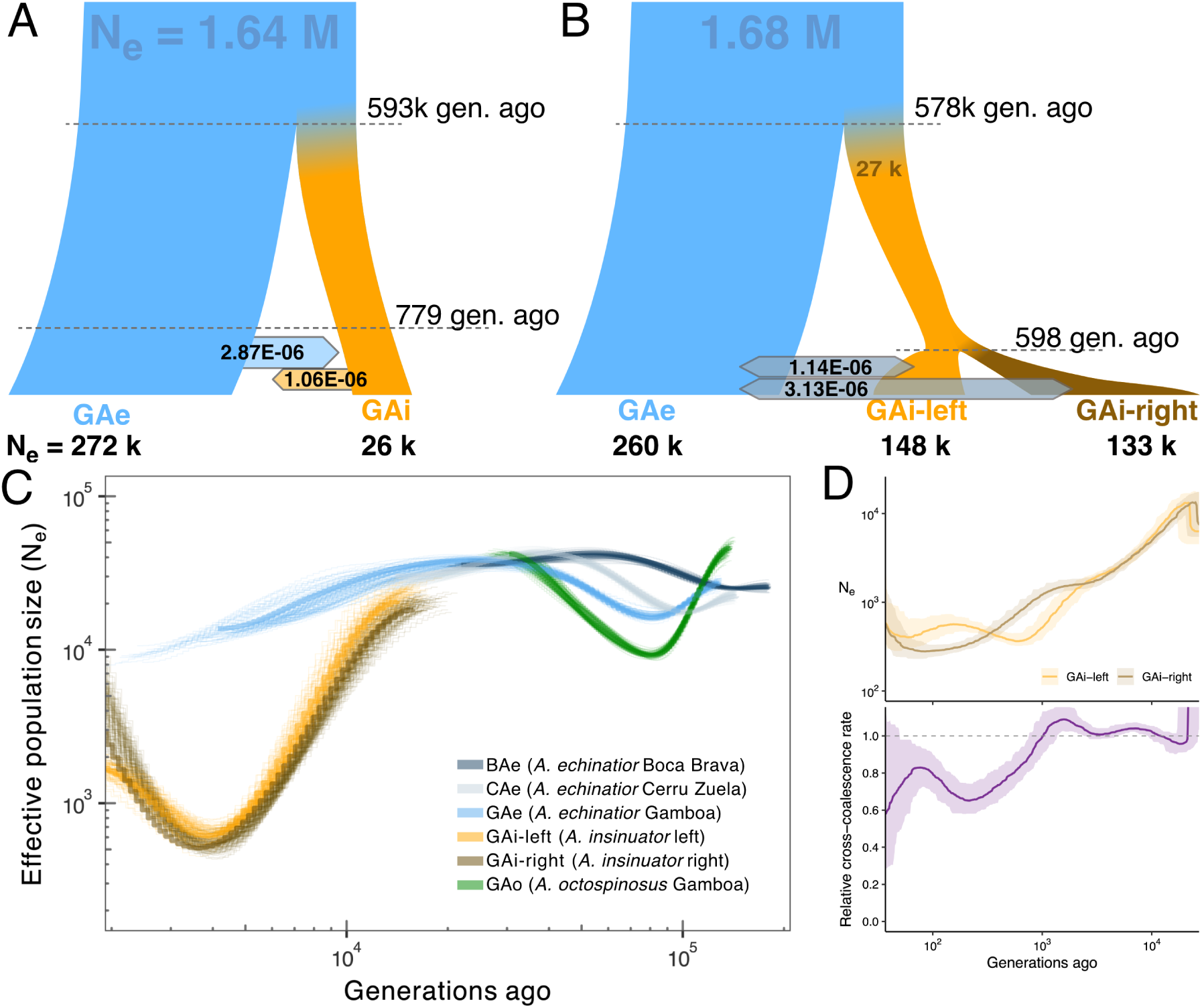
Demographic modelling of the *A. echinatior* and *A. insinuator* populations from Gamboa. **(A)** The best fitting two population model inferred that the split between the two species happened ∼600 k generations ago (95% CIs: 512606 -870638 generations), with the parasite lineage emerging as a fraction of the host population (N_e_ of *A. insinuator* = 25,636 (95% CIs: 19,274 – 33,754), N_e_ of *A. echinatior* = 270954 (95% CIs: 228573 - 344354). In that model, gene flow ceased after the speciation event but was resumed secondarily 779 generations ago (95% CIs: 415 - 1412), continuing into the present. See text for further statistics. **(B)** The best fitting three population model inferred the split between GAi-left and GAi-right to have occurred ∼600 generations ago and to have implied very limited reciprocal gene flow from the host *A. echinatior* into both *A. insinuator* demes ever since (GAe↔GAi-left: 1.1e-6, 95% CI: 0.8 - 4.7e-6; GAe↔GAi-right: 3.1e-6, 95% CI: 2.0 – 13.0e-6). (**C**) Effective population size changes between 2000 to 200000 generations ago as inferred with smc++ in *A. echninatior* from Gamboa, Boca Brava and Cerro Zuela, *A. octospinosus* from Gamboa and both *A. insinuator* demes from Gamboa. **(D)** Effective population size (top) and relative cross-coalescence (bottom) of the two GAi demes over the last 50 to 20,000 generations ago, inferred with MSMC2. Both demes show declining effective population sizes toward the present, while relative cross-coalescence begins to decrease from values close to 1 approximately 1,000 generations ago, consistent with progressive differentiation between the demes. Solid lines show median estimates across 134 genomic bootstraps and shaded ribbons indicate the 95% bootstrap intervals; the dashed horizontal line represents a putative rate of 1 (panmixia).

To accommodate the sub-structuring of GAi into two demes, we additionally fitted a set of three-population models (GAe, GAi-left and GAi-right), which recovered the same core demographic history (Figure 5B, Supplementary Figure 6): First, GAe and GAi diverged 576 k generations ago (95% CI: 506 k – 632 k). Second, GAe exceeded the GAi ancestor in effective population size by roughly an order of magnitude (GAe: 260 k, 95% CI: 229 k–282 k; GAi ancestor: 27 k, 95% CI: 21 k–35 k). Third, also this analysis recovered recent, low-level gene flow between the host and both parasite demes. As in the two-population analysis, the best-fitting three-population model restricted gene flow to the period following the GAi-left/GAi-right divergence, with no support for gene flow between GAe and the GAi ancestor, consistent with recent secondary contact. The two GAi subpopulations were inferred to have diverged very recently (600 generations, 95% CI: 146–948) and to have larger contemporary sizes than their common ancestor, though with wide confidence intervals in both cases (GAi-left: 148 k, 95% CI: 45 k–227 k; GAi-right: 133 k, 95% CI: 33 k–216 k). We did not model gene flow between the two GAi demes, as their very recent divergence would not allow separating effects of recent divergence and migration from earlier divergence and gene flow. However, given their F_ST_ value of 0.11, which is considerably below the divergence between BAe and GAe (F_ST_ = 0.38, Figure 4C) these two demes appear not to be reproductively isolated.

To further investigate historical changes in Nₑ, we applied smc++ to the populations of *A. echinatior*, *A. octospinosus* and *A. insinuator* (Figure 5C). Differences in genetic composition among the hosts and the social parasite affected the time window over which Nₑ could be reconstructed, spanning ∼10 k to 100 k generations before present in the host and ∼1 k to 10 k generations in the parasite. In *A. octospinosus* we identified a bottleneck ∼100,000 generations ago, coinciding with a more subtle decline in the *A. echinatior* populations from Gamboa and Cerro Zuela. Overall, *A. echinatior* populations remained largely stable over time. In the parasite, smc++ inferred a substantial bottleneck around 5,000 generations ago in both demes, after which the trajectories of GAi-left and GAi-right diverged; both demes then showed very recent expansions, consistent with the three-population model above. However, in these very recent times SMC++ is least constrained, and abrupt recent increases of this kind are a known limitation of the model. We used MSMC2 to further infer changes in effective population size and divergence for the two *A. insinuator* demes in very recent times (Figure 5D). These analyses indicated that N_e_ decreased by an order of magnitude over the last ∼20,000 generations, with a subtle increase in the very recent past (<100 generations ago). Relative cross-coalescence rate between the two demes started decreasing from 1 approximately 1000 generations ago, indicating progressive but incomplete reproductive separation, largely consistent with the results from the other demographic models (Figure 5B,C).

### Reduced efficiency of positive and purifying selection in social parasites

We explored to what extent the efficiency of natural selection differs between the socially parasitic and free-living *Acromyrmex* species, comparing allele frequency spectra and nucleotide diversity at 4-fold synonymous and 0-fold non-synonymous sites. As expected from the overall reduction of nucleotide diversity in the social parasite, π was reduced at coding sites by over an order of magnitude in *A. insinuator* compared to both free-living *Acromyrmex* species (Figure 6a). The ratio of nucleotide diversity at non-synonymous *versus* synonymous sites, π_N_/π_S_, was more than twice as high in *A. insinuator* as in any population of free-living *Acromyrmex* and clearly separated by bootstrap 95% CIs (Figure 6b).

**Figure 6:**
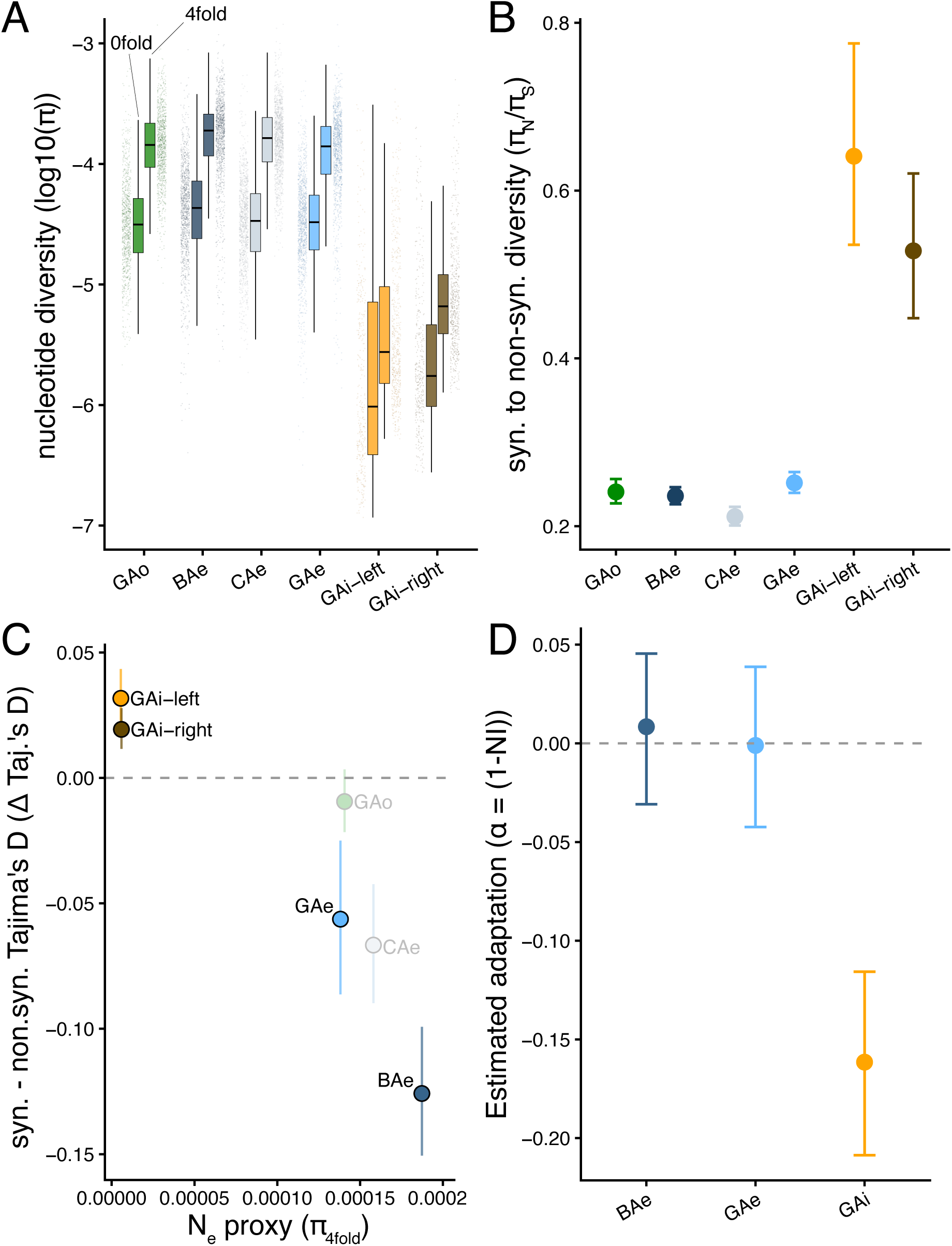
Signatures of reduced selection efficiency in *A. insinuator* relative to sympatric free-living *Acromyrmex* species. **(A)** Distribution of nucleotide diversity (log₁₀π) at 0-fold nonsynonymous and 4-fold synonymous sites across the genomes of the investigated (sub)populations of *A. octospinosus*, *A. echinatior* and *A. insinuator* calculated for 200 kb sliding windows. Boxes show median and interquartile ranges; jittered point ranges show the individual window distributions. **(B)** Genome-wide ratio of nonsynonymous to synonymous nucleotide diversity (π_N_/π_S_), computed by pooling per-site heterozygosities across all callable 0-fold and 4-fold sites. Error bars are 95% bootstrap confidence intervals from 2,000 replicate resampled windows. **(C)** The difference in Tajima’s D between 0-fold (nonsynonymous) and 4-fold (synonymous) sites (ΔTajima’s D) plotted against π_4fold_, a proxy for effective population size. Each point represents the genome-wide mean for one population; error bars are 95% bootstrap CIs from window resampling as in the previous panel. **(D)** Genome-wide proportion of nonsynonymous substitutions attributable to positive selection (α = 1 − (p_N_/p_S_)/(d_N_/d_S_)) for the three focal populations with sufficient sample size (BAe, GAe, GAi), computed using McDonald-Kreitman counts with *A. octospinosus* (GAo) as outgroup. Error bars are 95% bootstrap CIs from gene resampling (2,000 replicates). The strongly negative α in GAi reflects a substantial slightly-deleterious polymorphic load and highly constrained adaptive evolution (see also the load-corrected α estimates based on DFE-inference in the main text). Color coding throughout: GAo (green), BAe (dark blue), CAe (medium blue), GAe (light blue), GAi- left (orange) and GAi-right (brown).

These results, suggesting that the reduced effective population size of *A. insinuator* impairs the efficiency of natural selection, were corroborated by analyses of Tajima’s D at coding sites (Figure 6C). ΔTajima’s D, capturing the difference between nonsynonymous and synonymous sites, co-varied systematically with π_4fold_ (as a proxy for effective population size) across populations of the different species. Both *A. insinuator* clusters (GAi-left and GAi-right; Figure 2C) showed slightly positive ΔTajima’s D values, consistent with nonsynonymous variants segregating at similar frequencies as synonymous variants. In contrast, populations of free-living *Acromyrmex* all showed negative ΔTajima’s D values, consistent with purifying selection effectively reducing deleterious nonsynonymous variants to low frequencies. Note that the means for GAo and CAe fit the general trend but should be interpreted with caution given the small sample sizes (n = 3) for these populations.

We next estimated the genome-wide proportion of non-synonymous substitutions driven by positive selection (α = 1 - (p_N_/p_S_) / (d_N_/d_S_)) for GAe, BAe and GAi using McDonald-Kreitman statistics calculated with degenotate. Pairwise bootstrap comparisons indicated that the negative α of GAi (-0.16 [-0.21, -0.12]) differed significantly from the two very similar free-living primary host populations, which had estimated values close to zero: GAe (0.001 [-0.04, 0.04] and BAe (0.01 [-0.03, 0.05], p < 0.001 in both comparisons), whereas GAe and BAe did not differ significantly from each other (p = 0.73, Figure 6D). The substantially negative values for GAi indicate that α is systematically underestimated due to an excess of segregating slightly deleterious alleles at polymorphic loci, in line with a much lower effective population size, strong genetic drift and reduced efficiency of purifying-selection, as also inferred from our within-population diversity statistics (Figure 4, Figure 6A-C).

To resolve the deflation of α in GAi (Eyre-Walker and Keightley 2009; Messer and Petrov 2013), we complemented our analyses by inferring the distribution of fitness effects (DFE) in *A. insinuator* (GAi) and *A. echinatior* (GAe) using polyDFE v2.0 (Tataru et al. 2017). The unfolded site-frequency spectra showed an unexpectedly symmetric allele-frequency distribution at coding sites, suggesting substantial ancestral-state misassignment from the outgroup data. We therefore used folded SFS for DFE inference, despite having three diploid *A. octospinosus* samples available as outgroups. In both populations, Model C (deleterious + beneficial mutations) was preferred over Model B (deleterious mutations only), although the likelihood-ratio improvement of Model C over Model B was not statistically significant for GAi (LRT = 0.028, p = 0.99) and only marginally significant for GAe (LRT = 4.61, p = 0.10). However, model C is consistent with beneficial mutations occasionally contributing to adaptive parasite evolution, corresponding with the phenotypic studies summarized in the Introduction. It would also be consistent with the hypothesis that the two sympatric demes of *A. insinuator* in the Gamboa area are at least partly maintained by natural selection because they appear to have different host ant spectra.

The two species nonetheless yielded markedly different DFE inferences. The GAi population showed weak mean purifying selection (|S_d| = 0.050, with parametric estimates of 95% CIs derived from the polyDFE likelihood being 0.020 - 0.091), a minute fraction of beneficial mutations (p_b ≈ 6×10⁻⁹), and no detectable adaptive substitution after correcting for slightly-deleterious load (α_div = -0.007, 95% CI: -0.008 to -0.006). In contrast, the GAe population showed seven-fold stronger purifying selection (|S_d| = 0.340, 95% CI: 0.127 - 0.562) and a consistent but small fraction of beneficial mutations (p_b = 0.014%). We also inferred a significant fraction of (load-corrected) positive adaptive substitutions (α_div = 0.121, 95% CI: 0.018-0.215) in the free-living GAe population of *A. echinatior*, revealing a considerable load-bias in the raw McDonald-Kreitman estimates presented above. Together, these contrasting DFE inferences between the socially parasitic GAi population and the sympatric free living GAe population are consistent with fundamentally different evolutionary dynamics. This seems logical from the perspective that the social parasite evolved sympatrically from an ancestral population of its primary host species while adopting a fundamentally different life history.

## Discussion

The comparative resolution of our study, including whole genome data for more than 150 unrelated specimens from three different *Acromyrmex* species, provides unparalleled insights into the population genetics underlying the sympatric speciation history of an inquiline social parasite from its host (Figures 1-4). Our data also corroborated that *A. octospinosus* and *A. echinatior* split ca. 3.2 MYA (Vizueta et al. 2025), and that *A. echinatior* and *A. insinuator* diverged ∼1 MYA (Schrader et al. 2021). In addition, our data allowed powerful assessments of demography and interspecific geneflow (Figure 5), of differences in positive and purifying selection in the recent past, and of the adaptive potential of the specific gene pools considered (Figure 6). This was particularly relevant for the unexpected finding that the inquiline *A. insinuator* gene pool appears to be undergoing an incipient specialization process on *A. octospinosus* as alternative host, perhaps not unlike what the South American inquiline lineage of *A. charruanus* and *Pseudoatta* completed when these species split up <2 MYA (Schrader et al. 2021).

The *A. insinuator* inquiline parasite population that we studied in the Panama Canal Zone turned out to be dramatically genetically depleted, consistent with expectations from population genetics theory (Bourke and Franks 1991; Frankham et al. 2011) and from the general and *A. insinuator*-specific natural history records that we summarized in the Introduction. Even though *A. insinuator* expresses a number of clearly adaptive socially parasitic traits, the overall adaptive potential of its single known population appears severely constrained due to an overall reduction in the efficiency of natural selection. While our findings thus add to existing empirical and theoretical work ascribing a reduced capacity for adaptive evolution to inquiline social parasites like *A. insinuator* (Schrader et al. 2021), the question of how the substantial suite of adaptive phenotypic changes accompanying transitions from free-living colony-life to inquiline social parasitism can evolve remains unanswered. We hypothesize that the answer might ultimately relate to unusually strong positive selection for parasitic traits while the sympatric speciation process was ongoing until the parasitic deme became reproductively isolated by a very asymmetric form of disruptive selection, but further work will be needed to test this conjecture.

Although the separate species status of *A. insinuator* is not in doubt and the parasite’s sympatric emergence is well documented (Schultz et al. 1998; Schrader et al. 2021), we found clear signatures of limited, secondary gene flow between the extant Gamboa *A. insinuator* (GAi) population and the sympatric (GAe) and allopatric (CAe) host populations of *A. echinatior*, but presumably not with the more remote allopatric BAe population or the GAo population of *A. octospinosus*, the phylogenetically more remote secondary host for *A. insinuator* in Gamboa. This influx of host genetic variation into the population of the *A. insinuator* parasite could possibly alleviate the consequences of small effective population size on the efficiency of natural selection.

Our secondary contact demographic models (Figure 4) suggest that reproductive isolation between host and parasite became complete upon speciation ∼600 k generations ago but was secondarily relaxed in recent generations. Although we do not know the specific generation time of *A. insinuator* colonies, this re-establishment of limited and asymmetric gene flow must have been a Holocene event, i.e. it must have occurred less than ca. 11,700 years ago. The arrival of humans in Costa Rica and Panama is rather well documented to have started in the early Holocene (although scattered migration before the last Pleistocene ice age cannot be excluded), and farming human populations are known to have been well established more than 3000 years before present and to have had varying impact on their natural environment (Clement and Horn 2001). There is also good documentation for a natural mid-Holocene dry period in central America 5-8 kYA, which may have paved the way for increased human colonization (Olivares-Casillas et al. 2026). It thus seems possible that these natural and human disturbances of pristine rainforest habitat were instrumental in establishing larger and better-connected open habitat patches for species like *A. echinatior*, possibly allowing renewed contact between host and parasite populations that were not yet fully reproductively isolated.

Our present study provides ample support for reproductive isolation between Panamanian *A. echinatior* and *A. octospinosus*, justifying the distinct species status of *A. echinatior* as established by Schultz et al. (1998) based on segregation of a small set of diagnostic alleles. Our genome-wide analyses are thus conflicting with a recent study (Mera-Rodríguez et al. 2025) proposing to synonymize *A. echinatior* and *A. octospinosus* based on ultra-conserved element (UCE) data and variation in a single morphological trait (pronotal spine angle). While UCEs are a powerful approach to resolve deep phylogenetic splits, they tend to offer less resolution for shallow divergences (Meiklejohn et al. 2016). Furthermore, the original description of *A. insinuator* and *A. echinatior* by Schultz et al. (1998) made clear, based on a number of diagnostic allozyme alleles, that sympatric *A. echinatior* and *A. octospinosus* in Gamboa are reproductively isolated, a result that our present genome-wide SNP data confirm. Based on a survey of museum material from across Central and South America, Schultz et al. (1998) also concluded that the pronotal spine-angle trait works reasonably well for distinguishing between the sympatric species in Gamboa but becomes unreliable on its own outside Panama. They therefore offered additional morphological characters (e.g., pilosity or tubercle structure) which were not considered by Mera-Rodríguez et al. (2025). Their analyses also did not adjust for the substantial geographic covariation between UCE data and phylogenetic substructuring. Based on the data presented here, we thus recognize and reinstate the species *Acromyrmex echinatior*, Forel 1899, sp. rev., which was recently synonomized under *A. octospinosus* (Mera-Rodríguez et al. 2025).

### Population sub-structure and evolutionary dynamics of sympatric inquilines and hosts

By combining sampling-map data with genome-wide diversity information we were able to document a substantial degree of population viscosity in the *A. echinatior* population as it extended in our sampling from Gamboa village to Panama City. This finding helped to explain the even more substantial diversification between the GAe and the BAe populations several hundred km apart. We do not have comparable information for Panamanian *A. octospinosus* but would not be surprised if the populations of this species are less substructured because of its more continuous habitat. However, the long, species-level branches in the UCE-tree of Mera-Rodríguez et al. (2025) suggest that a number of cryptic species will ultimately be recognized within this *octospinosus*-complex, a diversification process that would be more likely to have occurred when population sub-structuring would be a general characteristic of Central American *Acromyrmex* ants.

Perhaps the most intriguing finding of our study is that the inquiline social parasite population in Gamboa is clearly sub-structured into two discrete lineages that originated in the Holocene, which suggests that similar cases of cryptic extant diversity may also characterize other inquiline social parasite lineages. We expect such secondary divergences to evolve more easily not only when populations are generally substructured but also when genomes are dramatically depleted and populations regularly remain so isolated that genetic drift becomes a major factor (Schrader et al. 2021). This could also contribute to the emergence of secondary host shifts followed by additional local adaptations, as observed in a convergently evolved inquiline parasite lineage of South American *Acromyrmex* that diverged from a free-living ancestor ∼5MYA and secondarily split <2 MYA into two species with distinct ecological and life history specializations (*A. ameliae* and *Pseudoatta argentina* (Schrader et al. 2021)).

In this context, it is remarkable that the two demes of *A. insinuator* in and around Gamboa had different host and temporal prevalences across the nearly 25 years of our sampling in Gamboa. In the 1990s, *A. insinuator* was almost exclusively collected from *A. echinatior* host colonies (GAi-right, the brown symbols and low nest numbers in Figure 3), while in the later years we started to increasingly find *A. insinuator* queens from what now turns out to be a genetically distinct cluster (GAi-left, orange, cf. Figure 2C) and in more equal or even opposite frequencies across *A. echinatior* and *A. octospinosus* host colonies. This turnover, and the increasing representation of *A. insinuator* queens obtained from *A. octospinosus* host colonies may be part of natural fluctuation dynamics, but we cannot exclude that our removal of large *A. echinatior* colonies with reproducing parasites from the GAi-right deme in the 1990s may have created open niche space for GAi-left parasites from *A. octospinosus* colonies in the forests surrounding Gamboa village. All this, and particularly our finding that the two demes of *A. insinuator* appear to exploit *A. echinatior* and *A. octospinosus* in different frequencies deserves further study.

## Data availability

Short read sequencing data for all 162 individuals are available in bioproject PRJNA1505145 on NCBI. Previously published data from six samples are available on NCBI under sample accessions SAMN18321427 (GAe-067), SAMN18321428 (GAe-077), SAMN18321430 (GAi-020), SAMN18321429 (GAi-073), SAMN18321431 (GAo-059), and SAMN18321432 (GAo-064). Code is available at https://zivgitlab.uni-muenster.de/AcroPopGen.

## Acknowledgements

The Visitor’s Office of the Smithsonian Tropical Research Institute provided logistic help and facilities to work in Gamboa and the Autoridad Nacional del Ambiente y el Mar (ANAM) gave permission to sample and export ants from Panama to Denmark throughout the fieldwork period. We are grateful to the dozens of researchers who have contributed to fieldwork in Panama to collect *Acromyrmex insinuator* and its two host species since 1993. We thank Volker Nehring and Ted Schultz for their helpful comments on the manuscript. This study was supported by grants from the European Research Council (ERC Advanced grant 323085) (J.J.B.) and the Lundbeck Foundation (R190-2014-2827) (G.Z.). Claude and github co-pilot were used for producing basic code, debugging and for condensing text in the initial draft of the manuscript. All AI-assisted text and code were reviewed and revised by the authors, who remain fully responsible for the content, analyses, and conclusions presented in this manuscript.

## Author Contributions

Conceptualization, L.S., J.J.B and G.Z.; Methodology: L.S., H.P., R.S.L., M.S., Q.L., G.Z.; Software: L.S., M.E., H.P.; Formal analysis: L.S., M.E., H.P., Q.L., J.J.B.; Investigation: L.S., J.J.B.; Resources: L.S., J.J.B., G.Z.; Data curation: L.S., J.J.B.; Writing – original draft: L.S., J.J.B; Writing – review & editing: All authors; Visualization: L.S., M.E.; Supervision: L.S., J.J.B., G.Z.; Project administration: L.S., J.J.B, G.Z.; Funding acquisition: J.J.B, G.Z.

## References

Alexander DH, Novembre J, Lange K. 2009. Fast model-based estimation of ancestry in unrelated individuals. Genome Res. 19:1655–1664.

Baer B. 2004. Male reproductive investment and queen mating-frequency in fungus-growing ants. Behav. Ecol. 15:426–432.

Barnett DW, Garrison EK, Quinlan AR, Strömberg MP, Marth GT. 2011. BamTools: a C++ API and toolkit for analyzing and managing BAM files. Bioinformatics 27:1691– 1692.

Bekkevold D, Boomsma JJ. 2000. Evolutionary transition to a semelparous life history in the socially parasitic ant acromyrmex insinuator. J. Evol. Biol. 13:615–623.

Boomsma JJ, Huszár DB, Pedersen JS. 2014. The evolution of multiqueen breeding in eusocial lineages with permanent physically differentiated castes. Anim. Behav. 92:241–252.

Boomsma JJ, Nash DR. 2014. Evolution: sympatric speciation the eusocial way. Curr. Biol. 24:R798–R800.

Borowiec ML, Cover SP, Rabeling C. 2021. The evolution of social parasitism in formica ants revealed by a global phylogeny. Proc. Natl. Acad. Sci. 118:e2026029118.

Bourke AFG, Franks NR. 1991. Alternative adaptations, sympatric speciation and the evolution of parasitic, inquiline ants. Biol. J. Linn. Soc. 43:157–178.

Buschinger A. 1986. Evolution of social parasitism in ants. Trends Ecol. Evol. 1:155–160.

Buschinger A. 1990. Sympatric speciation and radiative evolution of socially parasitic ants – heretic hypotheses and their factual background. Z. Zool. Syst. Evol. 28:241–260.

Buschinger A. 2009. Social parasitism among ants: a review (Hymenoptera: Formicidae). Myrmecological News:219–235.

Chang CC, Chow CC, Tellier LCAM, Vattikuti S, Purcell SM, Lee JJ. 2015. Second-generation PLINK: rising to the challenge of larger and richer datasets. GigaScience 4:7.

Clement RM, Horn SP. 2001. Pre-columbian land-use history in Costa Rica: a 3000-year record of forest clearance, agriculture and fires from laguna zoncho. Holocene 11:419–426.

Dahan RA, Rabeling C. 2022. Multi-queen breeding is associated with the origin of inquiline social parasitism in ants. Sci. Rep. 12:14680.

Danecek P, Auton A, Abecasis G, Albers CA, Banks E, DePristo MA, Handsaker RE, Lunter G, Marth GT, Sherry ST, et al. 2011. The variant call format and VCFtools. Bioinformatics 27:2156–2158.

Darwin C. 1859. On the origin of species by means of natural selection, or, the preservation of favoured races in the struggle for life. London: John Murray, Albemarle Street Available from: https://www.biodiversitylibrary.org/bibliography/82303

De Souza DJ, Soares IMF, Della Lucia TMC. 2007. Acromyrmex ameliae sp. n. (hymenoptera: formicidae): a new social parasite of leaf-cutting ants in Brazil. Insect Sci. 14:251–257.

Degueldre F, Mardulyn P, Kuhn A, Pinel A, Karaman C, Lebas C, Schifani E, Bračko G, Wagner HC, Kiran K, et al. 2021. Evolutionary history of inquiline social parasitism in plagiolepis ants. Mol. Phylogenet. Evol. 155:107016.

Delaneau O, Marchini J, Zagury J-F. 2012. A linear complexity phasing method for thousands of genomes. Nat. Methods 9:179–181.

D’Ettorre P, Heinze J. 2001. Sociobiology of slave-making ants. Acta Ethol. 3:67–82.

Errbii M, Gadau J, Becker K, Schrader L, Oettler J. 2024. Causes and consequences of a complex recombinational landscape in the ant *Cardiocondyla obscurior*. Genome Res. 34:863–876.

Errbii M, Keilwagen J, Hoff KJ, Steffen R, Altmüller J, Oettler J, Schrader L. 2021. Transposable elements and introgression introduce genetic variation in the invasive ant *Cardiocondyla obscurior*. Mol. Ecol. 30:6211–6228.

Ewels P, Magnusson M, Lundin S, Käller M. 2016. MultiQC: summarize analysis results for multiple tools and samples in a single report. Bioinformatics 32:3047–3048.

Excoffier L, Dupanloup I, Huerta-Sánchez E, Sousa VC, Foll M. 2013. Robust demographic inference from genomic and SNP data. PLOS Genet. 9:e1003905.

Eyre-Walker A, Keightley PD. 2009. Estimating the rate of adaptive molecular evolution in the presence of slightly deleterious mutations and population size change. Mol. Biol. Evol. 26:2097–2108.

Frankham R, Ballou JD, Briscoe DA. 2011. Introduction to conservation genetics. 2. ed., reprinted. Cambridge: Cambridge University Press

Gallardo A. 1916. Notes systématiques et éthologiques sur les fourmis attines de la république argentine. An. Mus. Nac. Hist. Nat. B. Aires 28:317–344.

Gray KW, Rabeling C. 2023. Global biogeography of ant social parasites: exploring patterns and mechanisms of an inverse latitudinal diversity gradient. J. Biogeogr. 50:316–329.

Hölldobler B, Wilson EO. 1990. The ants. Cambridge, Mass: Belknap Press of Harvard University Press

Howe J, Schiøtt M, Boomsma JJ. 2021. Queens of the inquiline social parasite acromyrmex insinuator can join nest-founding queens of its host, the leaf-cutting ant acromyrmex echinatior. Insectes Soc. 68:255–260.

Huxley J. 1930. Ants. J. Cape and H. Smith

Keightley PD, Jackson BC. 2018. Inferring the probability of the derived vs. the ancestral allelic state at a polymorphic site. Genetics 209:897–906.

Korunes KL, Samuk K. 2021. pixy: unbiased estimation of nucleotide diversity and divergence in the presence of missing data. Mol. Ecol. Resour. 21:1359–1368.

Kutter H. 1968. Die sozialparasitischen ameisen der schweiz. Neujahrsbl. Nat.forsch. Ges. Zür. 171:1–62.

Lambardi D, Dani FR, Turillazzi S, Boomsma JJ. 2007. Chemical mimicry in an incipient leaf-cutting ant social parasite. Behav. Ecol. Sociobiol. 61:843–851.

Lawson DJ, Hellenthal G, Myers S, Falush D. 2012. Inference of population structure using dense haplotype data. PLOS Genet. 8:e1002453.

Leppänen J, Seppä P, Vepsäläinen K, Savolainen R. 2016. Mating isolation between the ant myrmica rubra and its microgynous social parasite. Insectes Soc. 63:79–86.

Li H, Durbin R. 2009. Fast and accurate short read alignment with burrows–wheeler transform. Bioinformatics 25:1754–1760.

Liu H, Jia Y, Sun X, Tian D, Hurst LD, Yang S. 2017. Direct determination of the mutation rate in the bumblebee reveals evidence for weak recombination-associated mutation and an approximate rate constancy in insects. Mol. Biol. Evol. 34:119–130.

Meiklejohn KA, Faircloth BC, Glenn TC, Kimball RT, Braun EL. 2016. Analysis of a rapid evolutionary radiation using ultraconserved elements: evidence for a bias in some multispecies coalescent methods. Syst. Biol. 65:612–627.

Mera-Rodríguez D, Fernández-Marín H, Rabeling C. 2025. Phylogenomic approach to integrative taxonomy resolves a century-old taxonomic puzzle and the evolutionary history of the *Acromyrmex octospinosus* species complex. Syst. Entomol. 50:469–494.

Messer PW, Petrov DA. 2013. Population genomics of rapid adaptation by soft selective sweeps. Trends Ecol. Evol. 28:659–669.

Nehring V, Dani FR, Turillazzi S, Boomsma JJ, d’Ettorre P. 2015. Integration strategies of a leaf-cutting ant social parasite. Anim. Behav. 108:55–65.

Nehring V, Dijkstra MB, Sumner S, Hughes WOH, Boomsma JJ. 2018. Reconstructing the relatedness of cooperatively breeding queens in the panamanian leaf-cutting ant *Acromyrmex echinatior* (hymenoptera: formicidae). Myrmecol. News:33–45.

Okonechnikov K, Conesa A, García-Alcalde F. 2016. Qualimap 2: advanced multi-sample quality control for high-throughput sequencing data. Bioinformatics 32:292–294.

Olivares-Casillas G, Correa-Metrio A, Bernal JP, Franco-Gaviria JF, Zawisza E, Curtis JH, Escobar J. 2026. Holocene trends in precipitation seasonality in the northern tropical americas. Quat. Sci. Rev. 379:109914.

Ortius-Lechner D, Gertsch PJ, Boomsma JJK. 2000. Variable microsatellite loci for the leafcutter ant *Acromyrmex echinatior* and their applicability to related species. Molecular Ecology:114–116.

Poplin R, Ruano-Rubio V, DePristo MA, Fennell TJ, Carneiro MO, Van der Auwera GA, Kling DE, Gauthier LD, Levy-Moonshine A, Roazen D, et al. 2018. Scaling accurate genetic variant discovery to tens of thousands of samples.

Purcell S, Neale B, Todd-Brown K, Thomas L, Ferreira MAR, Bender D, Maller J, Sklar P, de Bakker PIW, Daly MJ, et al. 2007. PLINK: a tool set for whole-genome association and population-based linkage analyses. Am. J. Hum. Genet. 81:559–575.

Quinlan AR, Hall IM. 2010. BEDTools: a flexible suite of utilities for comparing genomic features. Bioinformatics 26:841–842.

Rabeling C, Messer S, Lacau S, do Nascimento IC, Bacci M Jr, Delabie JHC. 2019. Acromyrmex fowleri: a new inquiline social parasite species of leaf-cutting ants from South America, with a discussion of social parasite biogeography in the neotropical region. Insectes Soc. 66:435–451.

Rabeling C, Schultz TR, Bacci M Jr, Bollazzi M. 2015. Acromyrmex charruanus: a new inquiline social parasite species of leaf-cutting ants. Insectes Soc. 62:335–349.

Rabeling C, Schultz TR, Pierce NE, Bacci M Jr. 2014. A social parasite evolved reproductive isolation from its fungus-growing ant host in sympatry. Curr. Biol. 24:2047–2052.

Schrader L, Pan H, Bollazzi M, Schiøtt M, Larabee FJ, Bi X, Deng Y, Zhang G, Boomsma JJ, Rabeling C. 2021. Relaxed selection underlies genome erosion in socially parasitic ant species. Nat. Commun. 12:2918.

Schrader L, Rinke JL, Errbii M, Teresi SJ, Van Den Bos E, Xiong Z, Vizueta J, Boomsma JJ, Gadau J, Zhang G. 2026. Transposable elements as evolutionary catalysts of ant macrodiversity. Sci. Adv. 12:eaee0306.

Schultz TR, Bekkevold D, Boomsma JJ. 1998. Acromyrmex insinuator new species: an incipient social parasite of fungus-growing ants. Insectes Soc. 45:457–471.

Sirviö A, Gadau J, Rueppell O, Lamatsch D, Boomsma JJ, Pamilo P, Page RE. 2006. High recombination frequency creates genotypic diversity in colonies of the leaf-cutting ant acromyrmex echinatior. J. Evol. Biol. 19:1475–1485.

Smith NGC, Eyre-Walker A. 2002. Adaptive protein evolution in drosophila. Nature 415:1022–1024.

Sumner S, Hughes WOH, Boomsma JJ. 2003. Evidence for differential selection and potential adaptive evolution in the worker caste of an inquiline social parasite. Behav. Ecol. Sociobiol. 54:256–263.

Sumner S, Hughes WOH, Pedersen JS, Boomsma JJ. 2004. Ant parasite queens revert to mating singly. Nature 428:35–36.

Sumner S, Nash DR, Boomsma JJ. 2003. The adaptive significance of inquiline parasite workers. Proc. R. Soc. Lond. B 270:1315–1322.

Tataru P, Bataillon T. 2019. polyDFEv2.0: testing for invariance of the distribution of fitness effects within and across species. Bioinformatics 35:2868–2869.

Tataru P, Mollion M, Glémin S, Bataillon T. 2017. Inference of distribution of fitness effects and proportion of adaptive substitutions from polymorphism data. Genetics 207:1103– 1119.

Vizueta J, Xiong Z, Ding G, Larsen RS, Ran H, Gao Q, Stiller J, Dai W, Jiang W, Zhao J, et al. 2025. Adaptive radiation and social evolution of the ants. Cell 188:4828–4848.e25.

Weismann A. 1893. The all-sufficiency of natural selection: a reply to herbert spencer. Available from: https://books.google.de/books?hl=en&lr=&id=IaeD9xM_08wC&oi=fnd&pg=PA309&dq=Weismann+(1893)+The+All+Sufficiency+of+Natural+Selection&ots=Mx18BvtVd4&sig=Simzui3ggZwLX_fku42x_5wM9MM

Wheeler WM. 1910. Ants: their structure, development and behavior. New York: Columbia University Press

Wheeler WM. 1923. Social life among the insects. New York: Harcourt, Brace and Co.

Wheeler WM. 1928. The social insects: their origin and evolution. London / New York: Kegan Paul, Trench, Trubner & Co. / Harcourt, Brace and Co.

Wilson EO. 1971. The insect societies. Cambridge, Mass., Belknap Press of Harvard University Press Available from: http://archive.org/details/insectsocieties0000wils

Wolf JI, Seppä P. 2016. Queen size dimorphism in social insects. Insectes Soc. 63:25–38.

Zheng X, Levine D, Shen J, Gogarten SM, Laurie C, Weir BS. 2012. A high-performance computing toolset for relatedness and principal component analysis of SNP data. Bioinformatics 28:3326–3328.

