## Supplementary Material for "Population genomics of the inquiline social parasite *Acromyrmex insinuator* and its leaf-cutting ant hosts *A. echinatior* and *A. octospinosus* reveals cryptic differentiation and reduced efficiency of selection in the parasite"

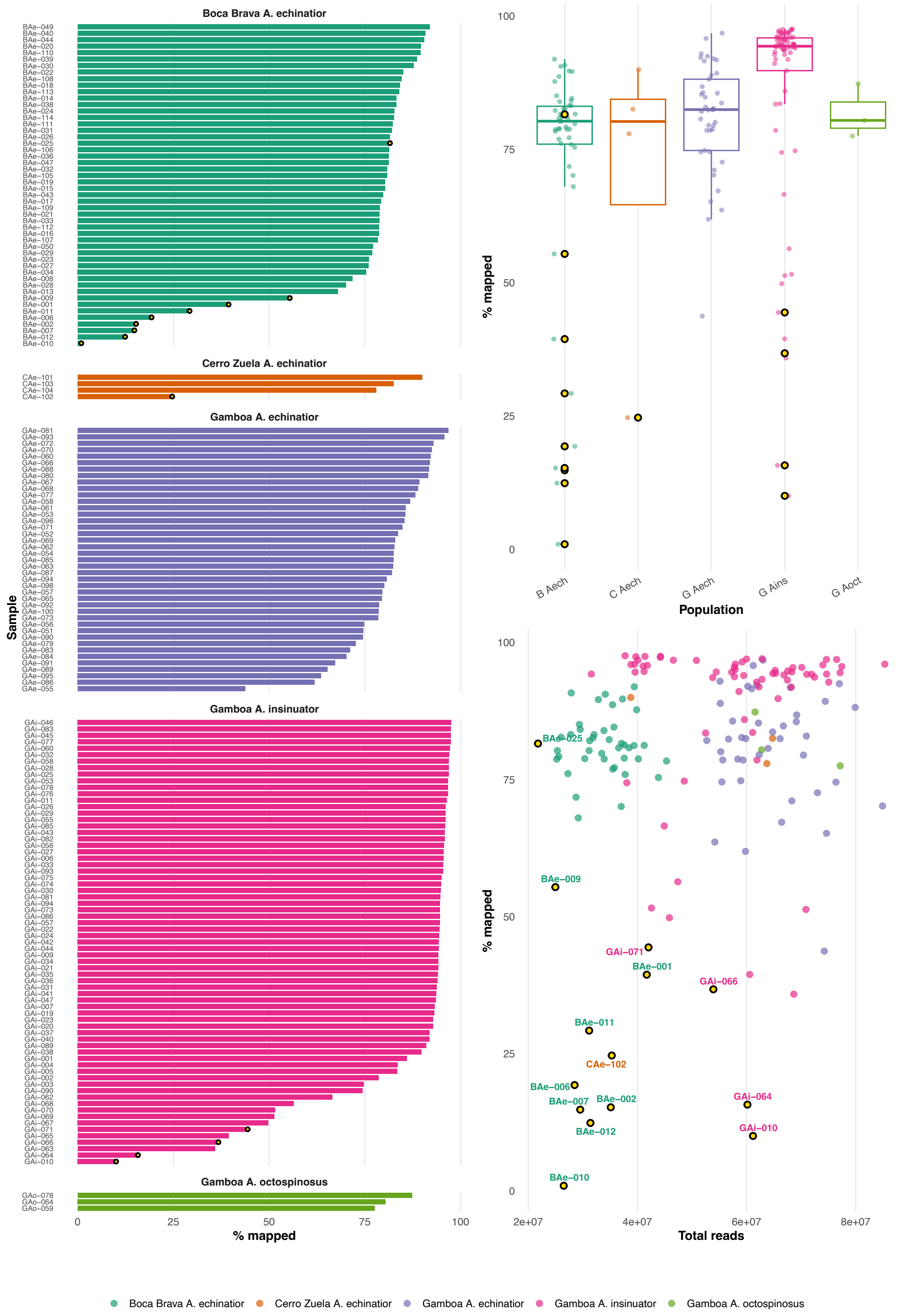


Supplementary Figure 1: Summary of short-read mapping rates and total number of reads generated per sample. Fourten samples excluded due to insufficient sequencing data quality are highlighted in yellow. Boxplots in the top right panel include data from the excluded samples. The final filtered dataset comprised 153 individuals (41 BAe, 3 CAe, 41 GAe, 65 GAi, and 3 GAo).

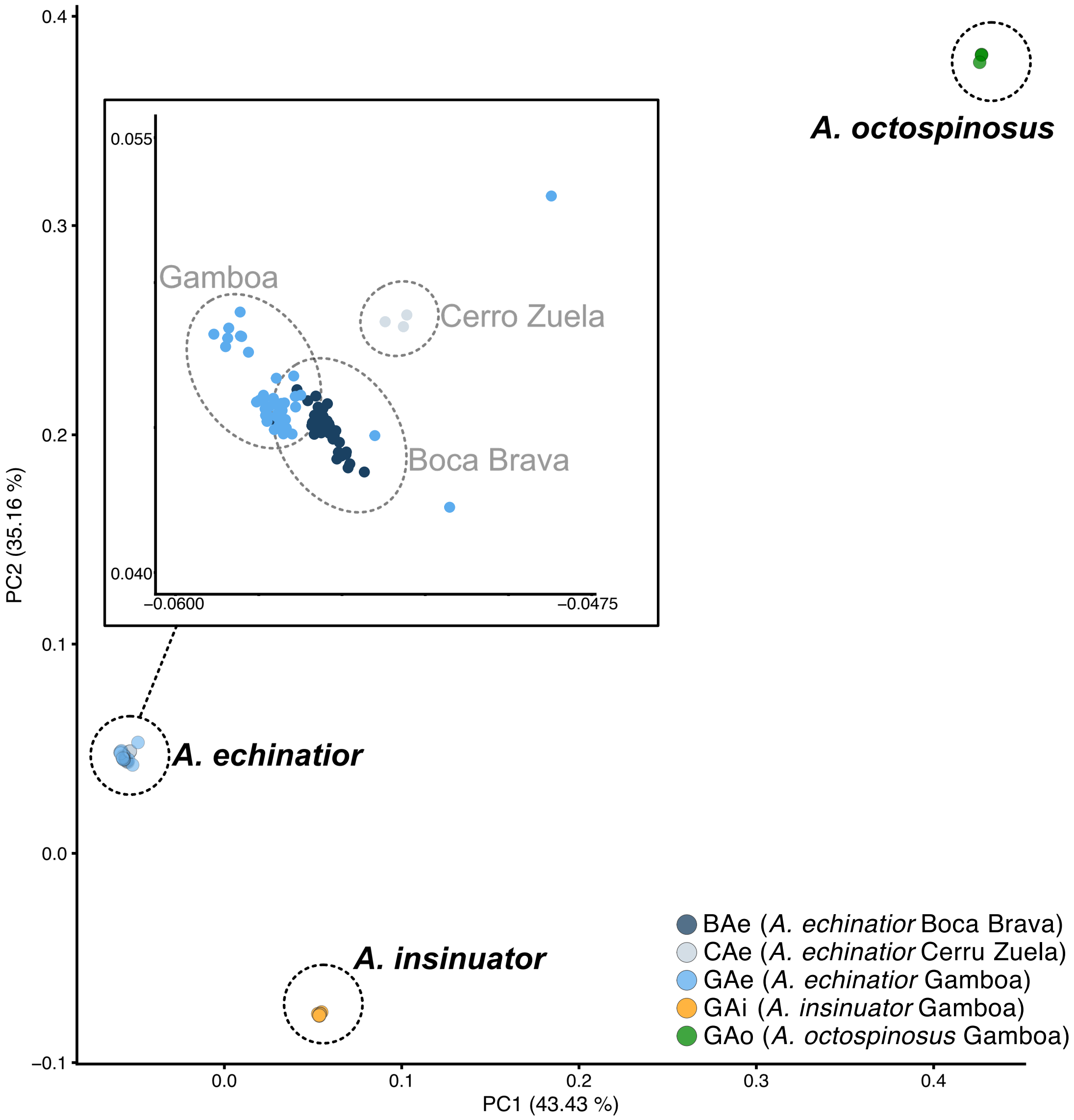


Supplementary Figure 2: Principal component analysis (PCA) across 138,179 LD-pruned, high-quality variants among 153 samples of *A. echinatior* from Boca Brava (dark blue) Gamboa (blue), and Cerro Zuela (light blue), *A. octospinosus* from Gamboa (green) and *A. insinuator* from Gamboa (orange). The insert in the top left is a zoom in on the *A. echinatior* cluster showing the separation of the three populations from Boca Brava, Gamboa and Cerro Zuela.

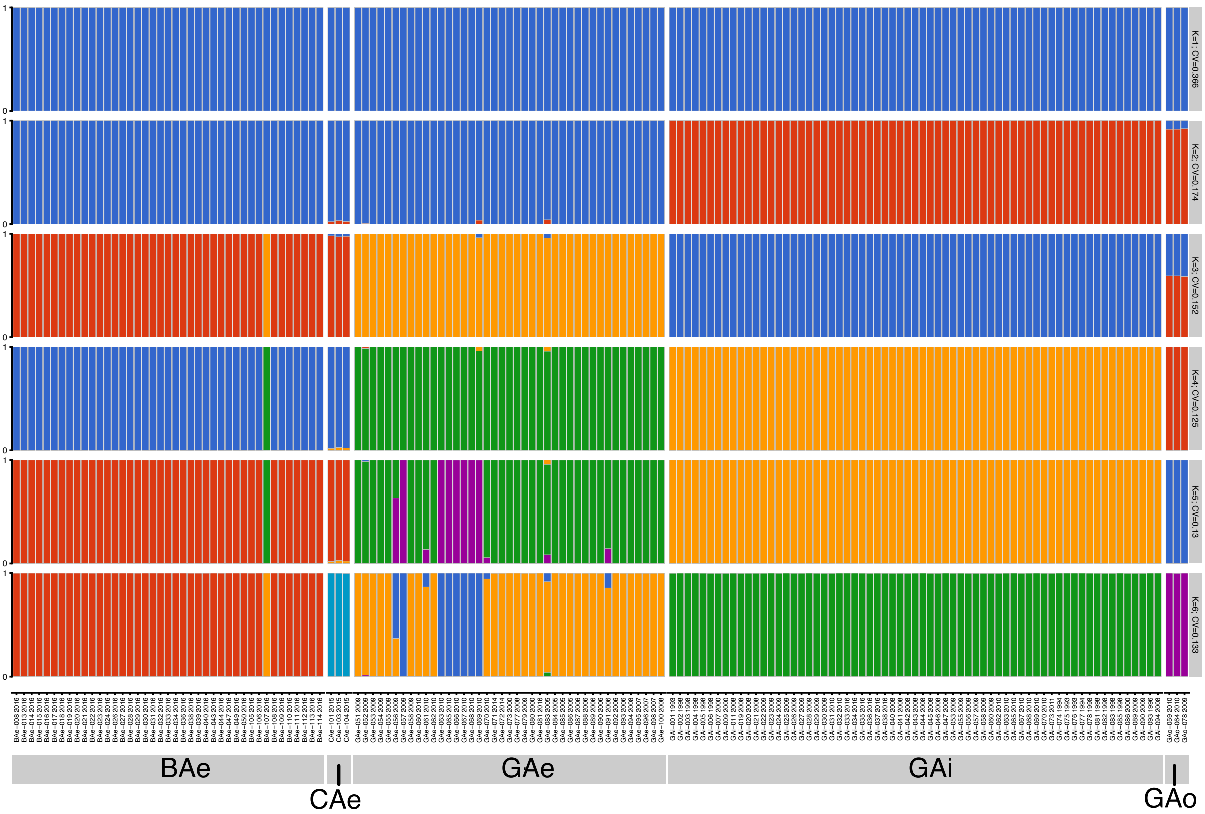


Supplementary Figure 3: ADMIXTURE analysis across 138,179 LD-pruned, high-quality variants among 153 of *A. echinatior*, *A. octospinosus* and *A. insinuator* for k=1 to k=6, with the best cross-validation (CV) error score reported for k=4 (CV=0.125).

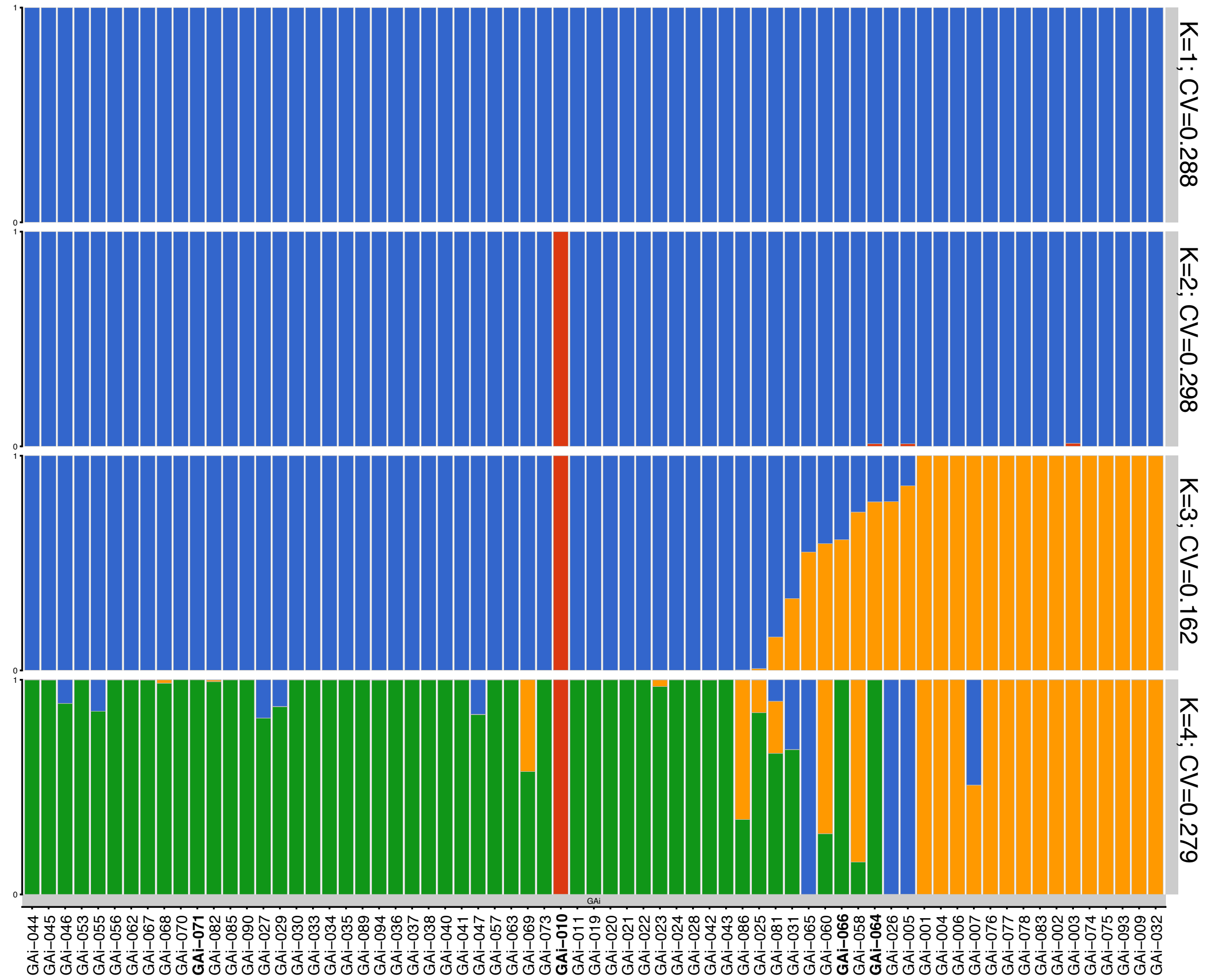


Supplementary Figure 4: ADMIXTURE analysis across LD-pruned, high-quality variants among 69 *A. insinuator* queens, including four samples that were removed from the general analyses due to insufficient sequencing quality (GAi-010, GAi-064, GAi-066, GAi-071, printed in bold) for k=1 to k=4, with the lowest cross-validation (CV) error score reported for k=3 (CV=0.162). The three clusters separate the two demes GAi-left (blue) and GAi-right (orange) and place GAi-010 in its own cluster. Note that GAi has the lowest total read count and the lowest mapping rate across all GAi samples. GAi-064 and GAi-066 are assigned to GAi-right and GAi-071 is assigned to GAi-left.

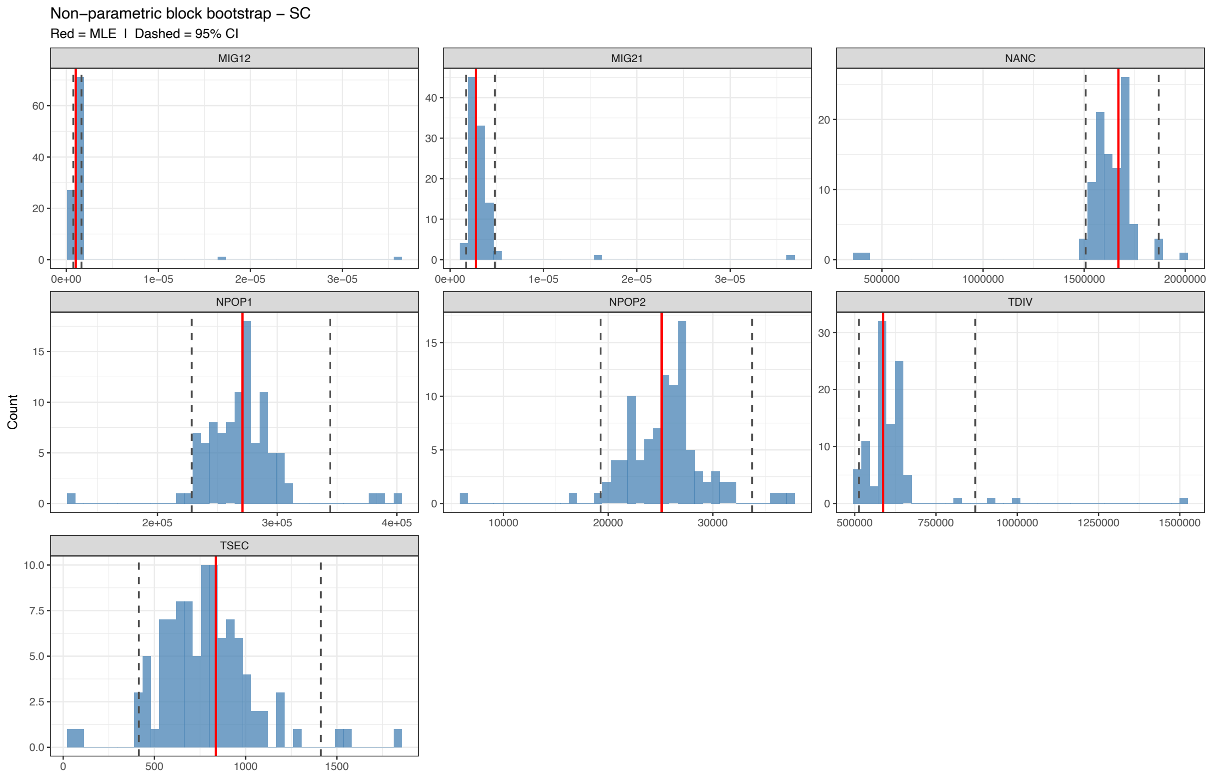


Supplementary Figure 5: Bootstrap distributions, maximum-likelihood estimates (MLE) and confidence intervals (CIs) derived from parametric block-bootstrapping of the best fitting demographic model (SC) from fastSimCoal2 of the separation between *A. insinuator* (GAi) and *A. echinatior* (GAe) from Gamboa, as shown in Figure 5A. MIG12= migration rate from GAe to GAi; MIG21 = migration rate from GAi to GAe; NANC = ancestral population size before the GAe-GAi split; NPOP1 = population size of GAe; NPOP2= population size of GAi; TDIV = GAe – GAi divergence time (in generations); TSEC = time of onset of secondary gene flow (in generations).

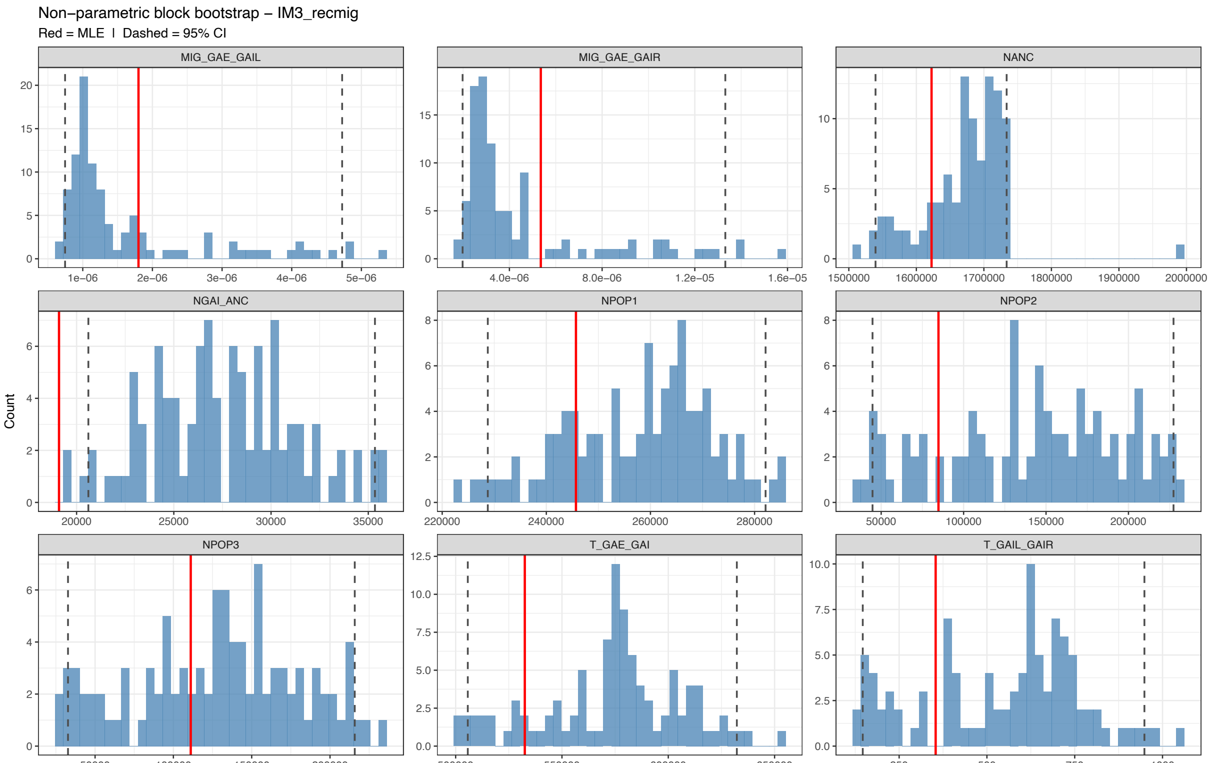


Supplementary Figure 6: Bootstrap distributions, maximum-likelihood estimates (MLE) and confidence intervals (CIs) derived from parametric block-bootstrapping of the best fitting demographic model (SC) from fastSimCoal2 of the separation between the two *A. insinuator* demes (GAi-left, GAi-right) and *A. echinatior* (GAe) from Gamboa, as shown in Figure 5B. MIG_GAE_GAIL = migration rate between GAe and GAi-left; MIG_GAE_GAIR = migration rate between GAe and GAi-right; NANC = ancestral population size before the GAe-GAi split; NANC_GAI = ancestral population size of GAi before the GAi-left – GAi-right split; NPOP1 = population size of GAe; NPOP2 = population size of GAi-left; NPOP3 = population size of GAi-right; T_GAE_GAI = GAe – GAi divergence time (in generations); T_GAIL_GAIR = GAi-left – GAi-right divergence time (in generations).
